# Mid-life and late life activities and their relationship with MRI measures of brain structure and functional connectivity in the UK Biobank cohort

**DOI:** 10.1101/2020.04.10.035451

**Authors:** Melis Anatürk, Sana Suri, Stephen M. Smith, Klaus P. Ebmeier, Claire E. Sexton

**Author notes:** Correponding Author: Melis Anatürk, Department of Psychiatry, University of Oxford, Warneford Hospital, Oxford, OX3 7JX.

## Abstract

**INTRODUCTION:** This study aimed to evaluate whether mid-life and late life participation in leisure activities is linked to measures of brain structure, functional connectivity and cognition in early old age.

**METHODS:** We examined data collected from 7,152 participants of the UK Biobank study. Weekly participation in six leisure activities was assessed twice. A cognitive battery and 3T MRI brain scan were administered at the second visit.

**RESULTS:** Weekly computer use at mid-life associated with larger volumes of the left putamen and higher scores for fluid intelligence, alphanumeric and numeric trail making tasks and prospective memory. Frequent attendance at a sports club or gym at mid-life was associated with stronger connectivity of the sensorimotor network with the lateral visual and cerebellar networks. No other associations were significant.

**DISCUSSION:** This study demonstrates that not all leisure activities contribute to cognitive health equally, nor is there one unifying neural signature across leisure activities.

## 1. Introduction

By 2050, the total number of older adults (i.e. individuals ≥ 60 years old) worldwide is expected to reach 2.1 billion [1]. While improvements in life expectancy is a significant achievement of the 21^st^ century, population ageing represents a major societal challenge [2]. This is because older adults are often at a greater risk of developing certain health conditions than their younger counterparts. For instance, an individual aged 90 or older has 25 times the risk of developing dementia compared to an individual in their late 60s [3,4]., Whole population rates of dementia, such as Alzheimer’s disease, are projected to triple by 2050 as a consequence of population ageing [1]. Consequently, there has been growing scientific interest in identifying modifiable factors that may reduce the risk of developing dementia and contribute to ‘better’ brain health in late life.

Leisure activities have been systematically linked to better cognitive performance and structural brain integrity in older adults [5–13]. However, one of the major drawbacks of published studies is the longstanding use of composite measures of leisure activities, which provides limited insight into the *specific* activities that should be targeted by interventions to promote brain health in older individuals. Targeting non-optimal activities may in part explain the limited efficacy of current randomized-controlled trials (RCTs) on cognitive and neural outcomes [14,15]. A small number of epidemiological studies have begun to probe more activity-specific associations, suggesting that not all activities equally contribute to the risk of cognitive impairment [16–18]. Importantly, a study that aims to evaluate whether different activities independently relate to markers of brain health requires a comparably larger number of univariate tests, relative to a neuroimaging study examining a single leisure activity score. With a sample that currently exceeds several thousand individuals, the UK Biobank study offers statistical power that allows a more fine-grained approach to examining the links between activities and the ageing brain.

The aim of this study was to investigate whether mid-life and late life leisure activities relate to MRI measures of grey matter (GM) volume, white matter (WM) microstructure, WM lesions, resting-state functional connectivity and cognitive function in late life. Based on previous findings [6,11,12], we predicted that more frequent participation in each of the activities assessed would correlate with better performance on tests of cognitive function. Prior work also informs the hypothesis that mid-life activities may have more consistent associations with cognition, relative to late life [19,20]. We further expected that more frequent leisure activity participation correlates with greater structural integrity, including greater regional GM and higher white matter integrity (i.e. higher fractional anisotropy (FA), lower mean diffusivity (MD) [10]). Given the limited evidence investigating activity-specific effects on resting-state functional connectivity, no predictions were made regarding this modality.

## 2. Methods

### 2.1 Sample characteristics

Data was provided by participants enrolled in the UK Biobank study, a large-scale prospective cohort study. These individuals were asked to complete a range of assessments including detailed lifestyle questionnaires, cognitive tests, physical measures (e.g. blood pressure and mobility tests), provide biological samples (e.g. blood, urine, saliva) and also provide permission to access their National Health Services (NHS) health records. Since 2014, a subsample of the original 500,000 participants have been invited back to undergo a single session of MRI scanning of the brain, body and heart, with the goal of reaching 100,000 scanned individuals by 2022. This sub-study is ongoing, with regular data releases made available to researchers [21]. At the time of paper preparation, imaging data from a total 15,000 participants had been released (January 2019).

In our analyses, we examine data collected from two phases: at recruitment (2006 – 2010) and MRI assessment (2014+; Supplementary Materials: Figure S1). The UK Biobank study received ethical approval from the NHS National Research Ethics Service (Ref 11/NW/0382) and all enrolled participants gave their informed and written consent.

### 2.2 Inclusion and exclusion criteria

The sample consisted of individuals without a diagnosis of stroke or dementia who had completed an MRI assessment and provided complete data on leisure activities and sociodemographic, health, cognitive and lifestyle variables (for flowchart, see Figure S2).

### 2.3 Activities

Activity levels were assessed through items delivered on a touch screen tablet. Weekly/less than weekly participation was examined for the following activities: going to a pub or social club; undertaking a religious activity; attending adult education classes; going to a sports club or gym; visiting friends and family and leisure-time computer use. Further information on these items is provided in the Supplementary Materials.

### 2.4 Cognitive function

A 15-minute study-specific battery of cognitive assessments was administered to participants via a touch screen tablet [22]. The cognitive measures examined are described in Table 1, with a detailed description in the supplementary materials.

**Table 1.**
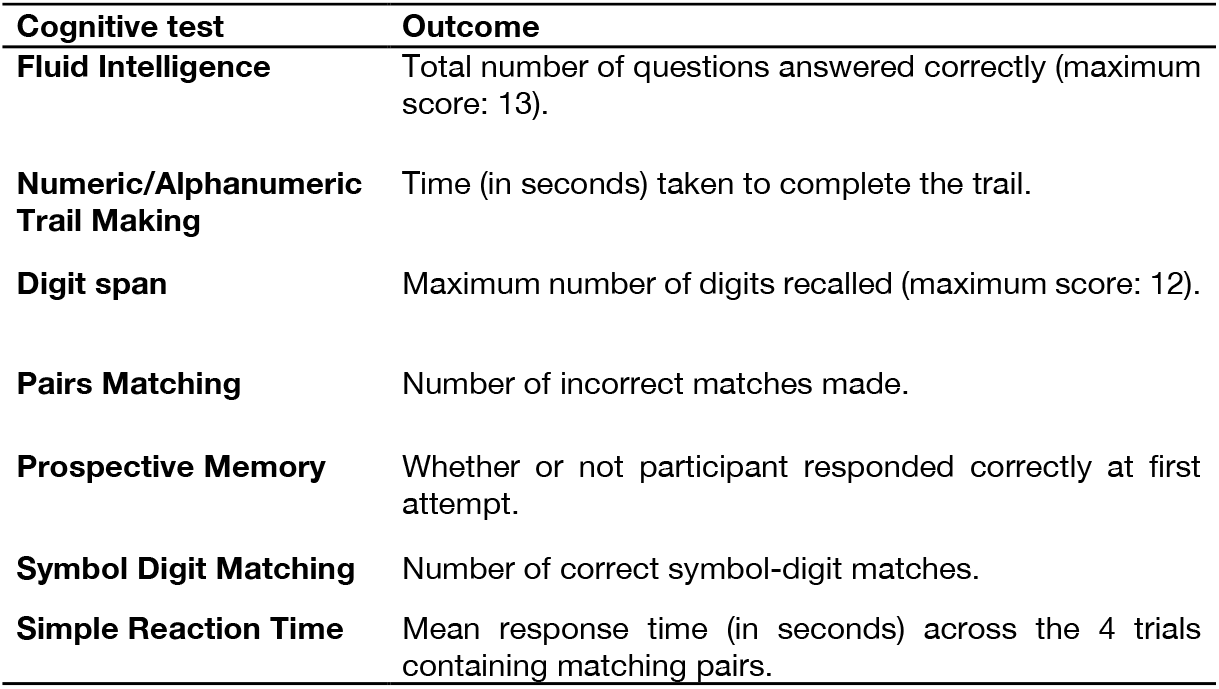
Neuropsychological tests of the UK Biobank battery examined in the present study.

### 2.5 Demographic and health-related variables

Baseline measures of age, sex, education, occupation, frequency of alcohol intake, sleep duration, body mass index (BMI), Mean arterial pressure (MAP) and social isolation (total number of individuals in household) were collected. ICD-10 diagnoses of depressive or anxiety disorders developed over the study duration was also examined. Additional information about these variables can be found in the Supplementary Materials.

### 2.6 MRI data acquisition and pre-processing

Participants were scanned using identical protocols with Siemens Skyra 3T (software VB13) and a Siemens 32-channel head coil at one of two study sites (i.e. Stockport or Newcastle).

T_1_-weighted images, diffusion-weighted images, T2 FLAIR images and restingstate functional images were assessed. Summary measures of brain structure and functional connectivity, or Image Derived Phenotypes (IDPs), have been generated on behalf of UK Biobank [23] and are available from UK Biobank upon data access application. For a detailed description of the imaging protocol and pre-processing steps, please see the supplementary materials.

### 2.7 Statistical analysis

All analyses were performed in R (version 3.5.2). The **lm** function in R (https://www.rdocumentation.org/packages/stats/versions/3.6.0/topics/lm) was used to fit a series of linear models to evaluate each activity as an independent predictor of the neuroimaging and cognitive measures. Specifically, the models assessed included each of the six activities, with confound co-variates including age, sex, education, occupational status, assessment centre, BMI, Mean Arterial Pressure (MAP), frequency of alcohol intake, sleep duration, the presence of depressive or anxiety disorders and the number of individuals living in a household. Separate models were run for midlife and later-life leisure activities, due to the possibility of autocorrelations between activity participation over time. Mean head motion and head size were also included as co-variates in the analysis of neuroimaging metrics. The imaging dependent variables consisted of total and regional GM volume (142 IDPs), total WM volume and lesions within WM (2 IDPs), WM microstructure (FA and MD in 27 pre-defined tracts, 54 IDPs), and partial correlation functional connectivity between large-scale resting-state networks (210 IDPs). A total of 8 cognitive outcomes were also assessed. Prospective memory was the only exception where an alternative to the linear regression was performed. Due to the binary nature of this variable (correct/incorrect answer), we used binomial logistic regression, where the outcome of interest was the log odds of completing the prospective memory task correctly at first attempt.

The distributions of all dependent variables were checked and in the case of non-normality, were log-transformed. Due to the number of univariate tests conducted, FDR-corrections were applied and are discussed here, with a full table of associations that were significant at a p_uncorrected_ < 0.05 level in Tables S2-S8. To facilitate these corrections, the **p.adjust** function in R was applied (https://www.rdocumentation.org/packages/podkat/versions/1.4.2/topics/p.adjust-methods), with FDR q values < 0.05 considered significant [24]. We report unstandardized beta-coefficients (B), their standard error (SE), standardized beta coefficients (β) and FDR q-values in the main text. Standardized beta coefficients were derived using the package **lm.beta** (https://cran.r-project.org/web/packages/lm.beta/lm.beta.pdf). For the logistic regression results, the odds ratio and 95% confidence intervals (95% C.I.) are reported.

## 3. Results

### 3.1 Participant demographics

A total of 7,152 participants were included in the analysis of neuroimaging outcomes. Participants were on average 56.39 years old (SD = 7.31) at baseline (i.e. “mid-life”) and 63.94 years old (SD = 7.32) at follow-up (i.e. “later-life”). Females represented 54.48% of the sample (n = 3,897). The percentage of participants reporting weekly participation at mid-life and later-life are listed in Table 2. A substantially smaller number of individuals had provided complete data for all cognitive measures at follow-up and were therefore analysed as a sub-set (n =1,734) of the main sample.

**Table 2.** Sample characteristics.

|  | **N (%) or Mean ±SD** | **Range** |
| --- | --- | --- |
| N. of participants | 7,152 |  |
| Duration between baseline and MRI scan (years) | 7.55 | 4.29 – 10.85 |
| **Demographics** |  |  |
| Age at baseline (years) | 56.39 ± 7.31 | 40 – 70 |
| Age at MRI scan (years) | 63.94 ± 7.32 | 46 – 80 |
| N. of females (%) | 3,897 (54.48) |  |
| Most common qualification earned (N.; %) | University/ college degree<br>(3,624; 50.67) |  |
| Most common occupational status (N.; %) | Professional occupations<br>(2,290; 32.02) |  |
| **Activities** |  |  |
| **N (%) reporting weekly participation in:** | **Mid-life** | **Later-life** |
| Leisure-time computer use | 6,340 (88.65) | 6,778 (94.77) |
| Visiting friends and family | 5,750 (80.40) | 5,874 (82.13) |
| Going to the pub or social club | 2,540 (35.51) | 2,513 (35.14) |
| Undertaking religious activities | 1,723 (24.09) | 1,770 (24.75) |
| Attending educational courses | 903 (12.63) | 775 (10.84) |
| Going to a sports club or gym | 3,852 (50.08) | 3,539 (49.48) |
| **Health and Lifestyle** |  |  |
|  | **N (%) or Mean ±SD** | **Range** |
| BMI (kg/m<sup>2</sup>) | 26.4 ± 4.01 | 16.14 – 55.07 |
| MAP | 99.57 ± 11.84 | 60.00 – 152.83 |
| Alcohol (frequency/week)* | 3.41 ± 1.35 | 0.00 – 5.00 |
| N (%) with ICD-10 diagnosis of Depression/Anxiety | 51 (0.71) |  |
| Sleep duration (hours/night) | 7.21 ± 0.93 | 2.00 – 13.00 |
| **Scanner** |  |  |
| N. of participants scanned at Stockport site (%) | 6,126 (85.65) |  |
| Relative head motion (mm) | 0.12 ± 0.06 | 0.03 – 1.39 |
| **Structural MRI measures** |  |  |
| Total GM (volume, mm<sup>3</sup>) | 613,301.98 ± 54,614.44 | 443,926 – 832,927 |
| WM (volume, mm<sup>3</sup>) | 547,421.16 ± 61,214.82 | 362,561 – 804,641 |
| CSF (volume, mm<sup>3</sup>) | 36,479.39 ± 17,032.15 | 7,613.27 – 157,075 |
| WM hyperintensities (volume, mm<sup>3</sup>) | 48,17.01 ± 6,150.75 | 30 – 86,534 |
| **Cognitive function (n = 1,734)** |  |  |
| Fluid intelligence score | 7.04 ± 1.93 | 1.00 – 13.00 |
| Alphanumeric trail making (seconds) | 52.28 ± 21.30 | 21.10 – 242.10 |
| Numeric trail making (seconds) | 21.5 ± 7.56 | 9.40 – 112.90 |
| Pairs matching test (total errors) | 6.52 ± 3.35 | 0.00 – 28.00 |
| Simple reaction time (seconds) | 0.59 ± 0.1 | 0.37 – 1.34 |
| N. of correct symbol-digit matches | 19.74 ± 5.14 | 2.00 – 36.00 |
| Backward digit span score | 6.91 ± 1.22 | 2.00 – 12.00 |
| Prospective memory score (N. % correct) | 1,510 (87.08%) |  |

**Note:** Values are Mean ± Standard deviation.
**Abbreviations-** BMI = Body mass index; BP = Blood pressure; CSF = Cerebrospinal fluid; GM = Grey matter; ICD = International Classification of Diseases; MAP = Mean arterial pressure; MRI = Magnetic resonance imaging; N = number; WM = White matter.

### 3.2 Cognitive function

Mid-life computer use was associated with better performance on the numeric (B = −0.04, SE = 0.009, β = −0.105, FDR q < 0.001) and alphanumeric (B = 0.032, SE = 0.01, β= −0.068, FDR q = 0.036) path trails (i.e. faster completion times) and a greater log odds of answering correctly on the prospective memory task (OR = 2.009, 95% C.I. = 1.336 – 22.971, FDR q = 0.014). Late life computer use was also associated with higher fluid intelligence scores (B = 0.701, SE = 0.2, β = −0.082, FDR q = 0.011) and faster completion of the alphanumeric (B = −0.049, SE = 0.014, β = −0.075, FDR q = 0.011) and numeric (B = −0.039, SE = 0.012, β = −0.073, FDR q = 0.015) trail making tasks and a higher log odds of correct responses on the prospective memory task (OR = 2.554, 95% C.I. = 1.507 – 4.208, FDR q = 0.011). Figure 1 presents an overview of the parallel associations between mid-life and late life computer use.

**Figure 1.**
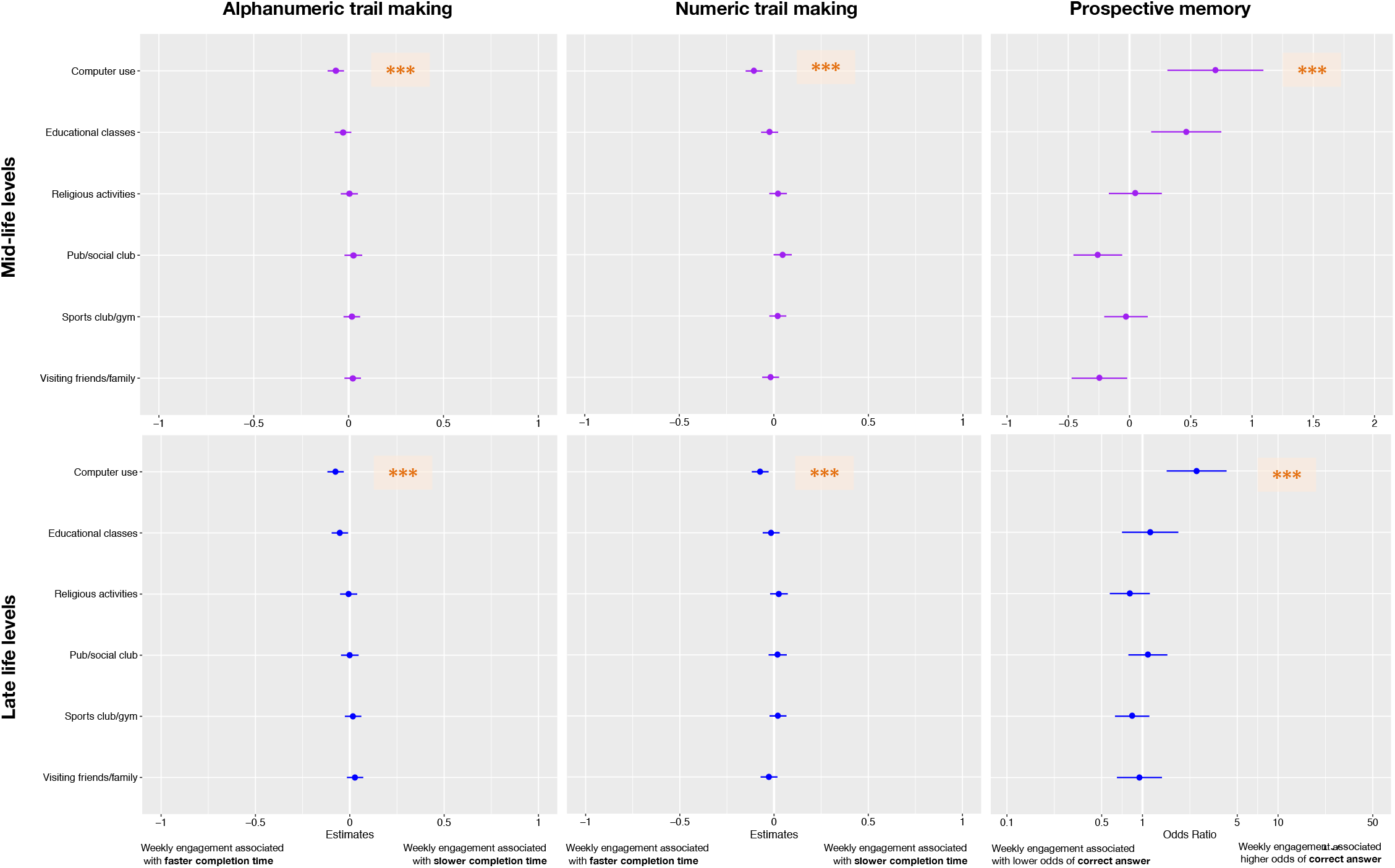
Dot-and-whisker plots to demonstrate associations between mid-life and late life computer use and alphanumeric trail making (standardized beta coefficients), numeric trail making (standardized beta coefficients) and prospective memory performance (odds ratio). Note: *** = survived FDR corrections. Adjusted for age, sex, education, occupational status, assessment centre, BMI, BP, frequency of alcohol intake, sleep duration, the presence of depressive or anxiety disorders, number of individuals living in a household and mutual activity effects.

Associations between late life attendance at educational courses and fluid intelligence use were also detected (B = 0.463, SE = 0.146, β = 0.075, FDR q = 0.015). No other activities, either during mid-life or later-life, were significantly linked to cognitive performance (FDR q’s > 0.05).

### 3.3 Structural MRI

Weekly computer use during mid-life, but not later-life was associated with greater volume measures in the left putamen (B = 77.429, SE = 0.17.534, β = 0.042, FDR q = 0.012). A trend in this direction was also observed for the right putamen, but it did not survive correction for multiple comparisons (Table S4). No significant associations were found for any other activity with GM volume, tract-specific FA or MD, total WM volume or lesions (FDR q’s > 0.05).

### 3.4 Functional MRI

Weekly attendance (during mid-life, but not late life) at a sports club or gym was associated with stronger absolute connectivity between the sensorimotor network and lateral visual network (B = 0.063, SE = 0.014, β = 0.056, FDR q = 0.009; Figure 2). Similarly, stronger connectivity was also observed between the sensorimotor network and cerebellar network for this activity (B = 0.062, SE = 0.014, β = 0.051, FDR q = 0.015; Figure 3). Post-hoc sensitivity analyses adjusting for total GM volume did not change the significance or effect size of the associations reported (data not shown).

There were no significant associations between any other activity and restingstate functional connectivity strength (FDR q > 0.05).

**Figure 2.**
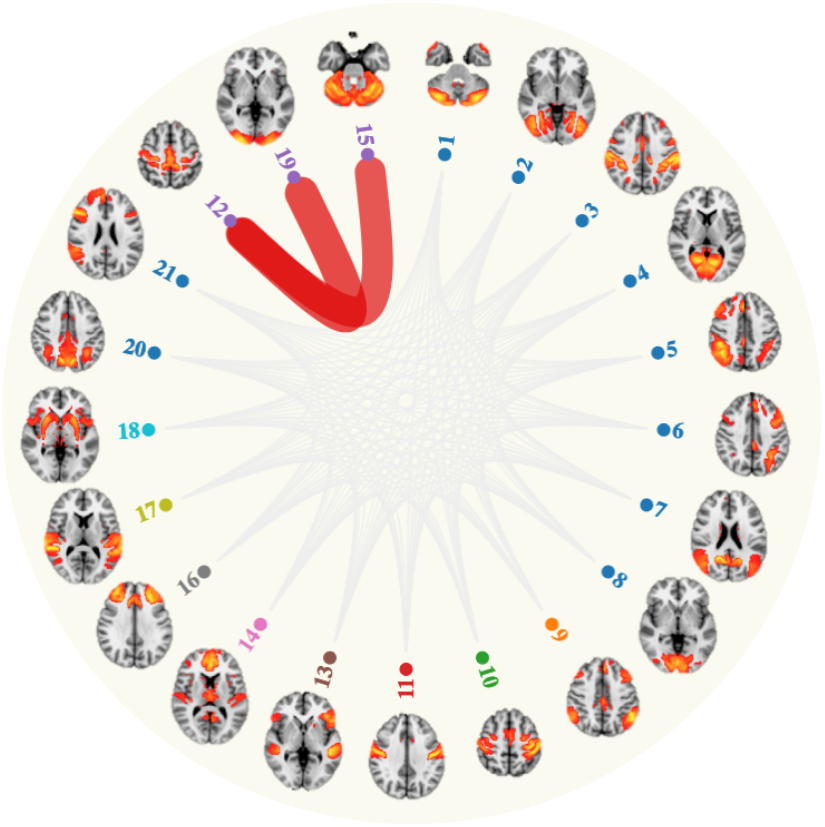
Mid-life sports club and gym attendance was associated with stronger connectivity (red lines) between the sensorimotor network (node 12) and lateral visual network (node 19) and cerebellar network (node 15). Results are adjusted for age, sex, education, relative motion, head size, occupational status, assessment centre, BMI, BP, frequency of alcohol intake, sleep duration, the presence of depressive or anxiety disorders, number of individuals living in a household and mutual activity effects. Associations reported are those that have survived FDR corrections for multiple comparisons.

## 4 Discussion

We present the largest multi-modal study to date to examine whether mid-life and late life activities are associated with brain integrity. We found that weekly leisure-time computer use was related to greater putamen volume and better performance on tests of cognition, relative to less frequent engagement. Furthermore, mid-life participation in a sports club or gym correlated positively with connectivity of the sensorimotor network, although demonstrated no associations with cognition. Attending an educational course in late life was also linked positively to fluid intelligence, but not with any of the MRI markers. We discuss each of these results in turn.

Individuals who used a computer for leisure on a weekly basis had more intact cognitive domains, including fluid intelligence, attention and task-switching, prospective memory and processing speed, following extensive co-variate adjustments. Parallel associations with cognitive test performance was also observed for late life computer use. These findings are in line with prior findings suggesting that frequent computer [25] and internet use [26] are linked to better cognitive function. Further, a large body of evidence support the notion that computerized cognitive training programmes lead to improvements in trained cognitive domains, with some findings suggesting transfer to untrained domains (e.g. [27,28]). Although speculative, computer use may be linked to cognition through an increased exposure to novelty (e.g. reading articles online), a greater demand on psychomotor skill (through mouse use and typing) and/or the opportunity to engage several domains of cognition at once (e.g. attention and memory when playing computer games; [29]. Our results may be taken to suggest that that interventions designed to encourage ‘active’ computer use could be administered to both middle-aged and older adults.

Despite the extensive links with cognition, the only region to be associated with mid-life computer use was GM volume in the left putamen. Given that this subcortical structure is involved in movement preparation and execution [30], it is perhaps expected that an activity requiring repeated use of the motor system (via typing and mouse use) might contribute to improvements in these abilities, as indicated by faster performance on the numeric and alphanumeric trail making tests. However, comparable improvements were not found in a separate task measuring simple reaction time, which suggests that a faster reaction time is unlikely to explain the correlations between computer use and performance on the trail making tasks. Notably, the putamen has more recently been implicated in non-motor functions, including executive control, working and episodic memory and category fluency [30]. The lateralization of results may be attributed to mouse use in a sample of predominantly right-handed individuals, although a trend observed between computer use and volumetric measures of right putamen suggests that mouse use may not be the only explanatory factor. Further, it is premature to conclude that computer use selectively contributes to putamen volume. For instance, a number of trends were observed with other measures of regional grey matter, tract-average FA and MD and functional connectivity (at a p_uncorrected_ < 0.05 level, see Tables S2-S8), prior to FDR corrections. These trends will, however, need to be independently validated by future studies before any conclusions can be drawn.

Another key finding was that attending a gym or sports club correlated with functional connectivity of several resting-state networks. Specifically, mid-life participation related to greater absolute connectivity of the sensorimotor network with the cerebellar and lateral visual network. Similar activation patterns at rest and during task execution indicate that both the sensorimotor and cerebellar networks may play a role in motor tasks [31], potentially interacting with one another to support motor behaviour. This is supported by the observation that subdivisions of the cerebellum are detected within the sensorimotor network [32,33], which was also observed in our study. Interestingly, the sensorimotor network further demonstrated stronger connectivity strength with the lateral visual network in individuals reporting weekly sports club/gym attendance, potentially suggesting enhanced visualmotor coupling. Different types of physical activities rely on visual information as a way of regulating and adapting current behaviour [34]. Our results are also in line with a study reporting increased connectivity between regions involved in the visual network and sensorimotor network, following a 13-week physical activity intervention [35].

Stronger connectivity *between* resting-state networks in an ageing sample is not entirely straightforward to interpret. For instance, older adults often exhibit less segregated sensorimotor networks (i.e. lower within-network connectivity and higher between-network connectivity) when compared to younger adults, with these increases in connectivity linked to poorer sensorimotor performance (e.g. reaction time, dexterity; [36]). Interestingly, we did not observe any neuropsychological correlates with this activity, although there was a trend for weekly gym/ sports club participation and faster reaction times (Table S2). Our findings, however, call for future investigations to evaluate whether increases in sensorimotor network connectivity associated with gym/sports club attendance relate to other behavioural markers (e.g. frailty, gait).

A final observation was that educational class attendance during later life correlated with higher fluid intelligence scores. These results resonate with a growing body of evidence in favour of life-long learning, and its contributions to maintaining cognitive health [5,37]. Notably, we did not identify parallel associations with mid-life educational class attendance. This may be taken to suggest that attending an educational class in adulthood may only bear transient effects on cognition, which places an importance on maintaining participation over time. However, as we only demonstrate a cross-sectional association, our study may simply be detecting a reverse causation: i.e. those who are more cognitively intact in late life choose to attend these educational courses. Results from the Baltimore Experience Corps trial provide evidence that enrolling older adults in an 30-hour intensive program (characterized by lectures, exercises, and interacting with children) based in elementary schools leads to improvements in cognition (i.e. flanker task performance; Carlson et al., 2015). We identified no associations with functional connectivity, GM or WM microstructure. This may be attributed to our examination of *weekly* course attendance, without the ability to distinguish between individuals who attend only once a week and to those who attend more frequently. Overall, our findings have practical implications, emphasizing the importance of educational courses designed for older adults. Notably, this was the least commonly reported activity in late life (11% of sample) which suggests that community-based initiatives to improve accessibility and participation levels in such programmes may be of value.

None of the other activities (i.e. visiting friends and family, going to the pub or social clubs, undertaking religious activities), measured in either mid-life or late life, were associated with markers of GM, WM microstructure, functional connectivity or cognitive function, after applying FDR corrections. While prior meta-analytic investigations have identified associations with brain structure [10] and cognition [39] when such activities are combined into composite scores, our results suggest that comparatively speaking, they do not uniquely contribute to brain health in early late life. Considering that the present findings imply dissociable effects between activities in brain-cognition associations, our results are in favour of an approach sensitive to these inter-activity differences, rather than the use of composite leisure activity scores, although the possibility has to be considered that cumulative activity over separate activities is required to improve brain function. Overall, our results are informative for both clinicians and researchers planning an RCT study, as they highlight several activities that may have an important role to play in maintaining neural and cognitive health among older adults.

### 4.1 Strengths and Limitations

The core strength of this study is the use of longitudinal data provided by a large cohort of middle-aged and older adults. The longitudinal design also provides insights into the neural and cognitive correlates of mid-life and late life activities, minimizing the risk of recall errors inherent in retrospective assessments of mid-life activity levels [20]. Furthermore, corrections for multiple comparisons were applied to minimize spurious findings, as larger studies are generally at a greater risk of identifying significant but clinically meaningless associations [21]. We were only able to examine six activities due to limited coverage in the Biobank study. Other common activities, such as reading [40] were not investigated. This study therefore represents an initial step towards better characterizing activity-specific associations with the brain but is by no means exhaustive. Further work is required to parse out the specific set of activities that have greater implications for brain ageing, generating evidence that may help improve current RCTs designs and retirement programmes in order to ensure that the most promising activities are targeted.

A major caveat of this work is the observational design of the study. Due to the opportunistic self-selected nature of the sample, we are unable to rule out reverse causation or residual confounding by a third unaccounted for variable. As an example of the latter, it may be the case that individuals who frequently use the computer are simply more accustomed to interacting with technology and therefore also perform better on the computerized tests of cognition. Cohort effects may alternatively explain our results. For example, while older individuals who are now fully engaged with technology may gain cognitive benefits, the same effects might not be observed in 20 years’ time as it becomes more common for individuals to become computer literate from a young age. We also note significant differences between individuals included in the sample to those excluded, with those included generally being older, more educated and from a higher occupational grade, being more likely to be female and differing in their lifestyle (e.g. alcohol, sleep duration) and health (BMI) parameters. These comparisons complement the observation that the larger Biobank cohort is generally healthier relative to the British general population (Fry et al., 2017). This would suggest that our results are most applicable to those who share similar characteristics to our sample and may not equally generalize to all middle- and older-aged adults.

## 5. Conclusion

This study identified computer use as associated with the left putamen volume and multiple domains of cognition. Furthermore, mid-life sports club/gym attendance was linked to sensorimotor connectivity, but not to cognitive performance. Educational classes in late life conversely held associations specific to fluid intelligence performance, with no other activities harbouring links with either cognitive domains, brain structure or functional connectivity. Overall, this study demonstrates selective associations between different leisure activities, highlighting several that may be relevant for RCTs aiming to promoting cognitive health in late life.

## 6. Acknowledgements

The authors thank all participants of the UK Biobank study who have dedicated their valuable time towards this project and the UK Biobank team for collecting, preparing and providing data used in this work. This research was conducted using the UK Biobank Resource under the approved application of 45301. The UK Biobank resource and brain imaging extension is funded by the UK Medical Research Council and the Wellcome Trust. M.A. is supported by the Clarendon Trust DPhil Fellowship and HDH Wills 1965 Charitable Trust (1117747). S.S. is supported by a fellowship from the UK Alzheimer’s Society (Ref 441), the EU Horizon 2020 Programme “Lifebrain” (Grant No. 732592) and WIN (203139/Z/16/Z). S.M.S. receives support from the Wellcome Trust (098369/Z/12/Z, 203139/Z/16/Z). K.P.E. reports support from the UK Medical Research Council (G1001354, MR/K013351/), the HDH Wills 1965 Charitable Trust (1117747), Alzheimer Research UK (PPG2012A-5) and the European Commission (Horizon 2020 grant “Lifebrain”, 732592). S.S. and C.E.S. are supported by the NIHR Oxford Biomedical Research Centre located at the Oxford University Hospitals NHS Trust and the University of Oxford, the NIHR Oxford Health BRC. The Wellcome Centre for Integrative Neuroimaging is supported by core funding from the Wellcome Trust (203139/Z/16/Z).

## Author contributions

S.M.S. provided the overall scientific strategy for UK Biobank brain imaging. M.A. planned and conducted the analyses and prepared the manuscript, including all tables and figures. S.S., S.M.S, C.E.S. and K.P.E. provided feedback and comments on all versions of the manuscript.

## References

[1] Prince MM, Wimo A, Guerchet M, Gemma-Claire A, Wu Y-T, Prina M, et al. World Alzheimer Report 2015 The Global Impact of Dementia An Analysis of Prevalence, Incidence, Cost and Trends. London: 2015.

[2] World Health Organization. World Report on Ageing and Health. Luxembourg: World Health Organization; 2015. https://www.who.int/ageing/events/world-report-2015-launch/en/.

[3] Yip AG, Brayne C, Matthews FE, MRC Cognitive Function and Ageing Study. Risk factors for incident dementia in England and Wales: The Medical Research Council Cognitive Function and Ageing Study. A population-based nested case–control study. Age Ageing 2006;35:154–60. https://doi.org/10.1093/ageing/afj030.

[4] Brayne C. The elephant in the room — healthy brains in later life, epidemiology and public health. Nat Rev Neurosci 2007;8:233–9. https://doi.org/10.1038/nrn2091.

[5] Matyas N, Keser Aschenberger F, Wagner G, Teufer B, Auer S, Gisinger C, et al. Continuing education for the prevention of mild cognitive impairment and Alzheimer’s-type dementia: A systematic review and overview of systematic reviews. BMJ Open 2019;9:e027719. https://doi.org/10.1136/bmjopen-2018-027719.

[6] Wang H-XX, Xu W, Pei J-JJ. Leisure activities, cognition and dementia. Biochim Biophys Acta – Mol Basis Dis 2012;1822:482–91. https://doi.org/10.1016/j.bbadis.2011.09.002.

[7] Wassenaar TM, Yaffe K, van der Werf YD, Sexton CE. Associations between modifiable risk factors and white matter of the aging brain: insights from diffusion tensor imaging studies. Neurobiol Aging 2019;80:56–70. https://doi.org/10.1016/J.NEUROBIOLAGING.2019.04.006.

[8] Sexton CE, Mackay CE, Ebmeier KP. A Systematic Review and Meta-Analysis of Magnetic Resonance Imaging Studies in Late-Life Depression. Am J Geriatr Psychiatry 2013;21:184–95. https://doi.org/10.1016/j.jagp.2012.10.019.

[9] Fratiglioni L, Paillard-Borg S, Winblad B. An active and socially integrated lifestyle in late life might protect against dementia. Lancet Neurol 2004;3:343–53. https://doi.org/10.1016/S1474-4422(04)00767-7.

[10] Anatürk M, Demnitz N, Ebmeier KP, Sexton CE. A systematic review and meta-analysis of structural magnetic resonance imaging studies investigating cognitive and social activity levels in older adults. Neurosci Biobehav Rev 2018. https://doi.org/10.1016/J.NEUBIOREV.2018.06.012.

[11] Evans IEM, Martyr A, Collins R, Brayne C, Clare L. Social Isolation and Cognitive Function in Later Life: A Systematic Review and Meta-Analysis. J Alzheimer’s Dis 2018;Preprint:1–26. https://doi.org/10.3233/JAD-180501.

[12] Yates LA, Ziser S, Spector A, Orrell M. Cognitive leisure activities and future risk of cognitive impairment and dementia: systematic review and meta-analysis. Int Psychogeriatr 2016;28:1–16. https://doi.org/10.1017/S1041610216001137.

[13] Erickson KI, Leckie RL, Weinstein AM. Physical activity, fitness, and gray matter volume. Neurobiol Aging 2014;35:S20–8. https://doi.org/10.1016/j.neurobiolaging.2014.03.034.

[14] Stephen R, Liu Y, Ngandu T, Antikainen R, Hulkkonen J, Koikkalainen J, et al. Brain volumes and cortical thickness on MRI in the Finnish Geriatric Intervention Study to Prevent Cognitive Impairment and Disability (FINGER). Alzheimers Res Ther 2019;11:53. https://doi.org/10.1186/s13195-019-0506-z.

[15] Mortimer JA, Ding D, Borenstein AR, DeCarli C, Guo Q, Wu Y, et al. Changes in brain volume and cognition in a randomized trial of exercise and social interaction in a community-based sample of nondemented Chinese elders. J Alzheimers Dis 2012;30:757–66. https://doi.org/10.3233/JAD-2012-120079.

[16] Krell-Roesch J, Syrjanen JA, Vassilaki M, Machulda MM, Mielke MM, Knopman DS, et al. Quantity and quality of mental activities and the risk of incident mild cognitive impairment. Neurology 2019;93:e548–58. https://doi.org/10.1212/WNL.0000000000007897.

[17] Fancourt D, Steptoe A, Cadar D. Cultural engagement and cognitive reserve: museum attendance and dementia incidence over a 10-year period. Br J Psychiatry 2018;213:661–3. https://doi.org/10.1192/bjp.2018.129.

[18] Krell-Roesch J, Vemuri P, Pink A, Roberts RO, Stokin GB, Mielke MM, et al. Association Between Mentally Stimulating Activities in Late Life and the Outcome of Incident Mild Cognitive Impairment, With an Analysis of the *APOE* ε4 Genotype. JAMA Neurol 2017;74:332. https://doi.org/10.1001/jamaneurol.2016.3822.

[19] Chan D, Shafto M, Kievit R, Matthews F, Spink M, Valenzuela M, et al. Lifestyle activities in mid-life contribute to cognitive reserve in late-life, independent of education, occupation, and late-life activities. Neurobiol Aging 2018;70:180–3. https://doi.org/10.1016/J.NEUROBIOLAGING.2018.06.012.

[20] Gow AJ, Pattie A, Deary IJ. Lifecourse Activity Participation From Early, Mid, and Later Adulthood as Determinants of Cognitive Aging: The Lothian Birth Cohort 1921. J Gerontol B Psychol Sci Soc Sci 2017;72:25–37. https://doi.org/10.1093/geronb/gbw124.

[21] Miller KL, Alfaro-Almagro F, Bangerter NK, Thomas DL, Yacoub E, Xu J, et al. Multimodal population brain imaging in the UK Biobank prospective epidemiological study. Nat Neurosci 2016;19:1523–36. https://doi.org/10.1038/nn.4393.

[22] Cornelis MC, Wang Y, Holland T, Agarwal P, Weintraub S, Morris MC. Age and cognitive decline in the UK Biobank. PLoS One 2019;14:e0213948. https://doi.org/10.1371/journal.pone.0213948.

[23] Alfaro-Almagro F, Jenkinson M, Bangerter NK, Andersson JLR, Griffanti L, Douaud G, et al. Image processing and Quality Control for the first 10,000 brain imaging datasets from UK Biobank. Neuroimage 2018;166:400–24. https://doi.org/10.1016/J.NEUROIMAGE.2017.10.034.

[24] Cox SR, Lyall DM, Ritchie SJ, Bastin ME, Harris MA, Buchanan CR, et al. Associations between vascular risk factors and brain MRI indices in UK Biobank. Eur Heart J 2019;40:2290–300. https://doi.org/10.1093/eurheartj/ehz100.

[25] Tun PA, Lachman ME. The Association Between Computer Use and Cognition Across Adulthood: Use it so You Won’t Lose it? Psychol Aging 2010;25:560. https://doi.org/10.1037/A0019543.

[26] Berner J, Comijs H, Elmståhl S, Welmer A-K, Sanmartin Berglund J, Anderberg P, et al. Maintaining cognitive function with internet use: a two-country, six-year longitudinal study. Int Psychogeriatrics 2019;31:929–36. https://doi.org/10.1017/S1041610219000668.

[27] Kueider AM, Parisi JM, Gross AL, Rebok GW. Computerized Cognitive Training with Older Adults: A Systematic Review. PLoS One 2012;7:e40588. https://doi.org/10.1371/journal.pone.0040588.

[28] Nguyen L, Murphy K, Andrews G. Immediate and long-term efficacy of executive functions cognitive training in older adults: A systematic review and meta-analysis. Psychol Bull 2019. https://doi.org/10.1037/bul0000196.

[29] Tun PA, Lachman ME. The association between computer use and cognition across adulthood: Use it so you won’t lose it? Psychol Aging 2010;25:560–8. https://doi.org/10.1037/a0019543.

[30] Ell S, Helie S, Hutchinson S. Contributions of the putamen to cognitive function. In: Andres Costa, Eugenio Villalba, editors. Horizons Neurosci. Res. 7th ed., Nova Science Publishers Inc; 2011, p. 29–52.

[31] Smith SM, Fox PT, Miller KL, Glahn DC, Fox PM, Mackay CE, et al. Correspondence of the brain’s functional architecture during activation and rest. Proc Natl Acad Sci U S A 2009;106:13040–5. https://doi.org/10.1073/pnas.0905267106.

[32] Habas C, Kamdar N, Nguyen D, Prater K, Beckmann CF, Menon V, et al. Distinct cerebellar contributions to intrinsic connectivity networks. J Neurosci 2009;29:8586–94. https://doi.org/10.1523/JNEUROSCI.1868-09.2009.

[33] Dobromyslin VI, Salat DH, Fortier CB, Leritz EC, Beckmann CF, Milberg WP, et al. Distinct functional networks within the cerebellum and their relation to cortical systems assessed with independent component analysis. Neuroimage 2012;60:2073–85. https://doi.org/10.1016/j.neuroimage.2012.01.139.

[34] Mallek M, Benguigui N, Dicks M, Thouvarecq R. Sport expertise in perception-action coupling revealed in a visuomotor tracking task. Eur J Sport Sci 2017;17:1270–8. https://doi.org/10.1080/17461391.2017.1375014.

[35] Demirakca T, Cardinale V, Dehn S, Ruf M, Ende G. The Exercising Brain: Changes in Functional Connectivity Induced by an Integrated Multimodal Cognitive and Whole-Body Coordination Training. Neural Plast 2016;2016:8240894. https://doi.org/10.1155/2016/8240894.

[36] Cassady K, Gagnon H, Lalwani P, Simmonite M, Foerster B, Park D, et al. Sensorimotor network segregation declines with age and is linked to GABA and to sensorimotor performance. Neuroimage 2019;186:234–44. https://doi.org/10.1016/j.neuroimage.2018.11.008.

[37] Hatch SL, Feinstein L, Link BG, Wadsworth MEJ, Richards M. The continuing benefits of education: adult education and midlife cognitive ability in the British 1946 birth cohort. J Gerontol B Psychol Sci Soc Sci 2007;62:S404–14. https://doi.org/10.1093/geronb/62.6.s404.

[38] Carlson MC, Kuo JH, Chuang Y-F, Varma VR, Harris G, Albert MS, et al. Impact of the Baltimore Experience Corps Trial on cortical and hippocampal volumes. Alzheimer’s Dement 2015;11:1340–8.

[39] Kuiper JS, Zuidersma M, Zuidema SU, Burgerhof JGM, Stolk RP, Oude Voshaar RC, et al. Social relationships and cognitive decline: a systematic review and meta-analysis of longitudinal cohort studies. Int J Epidemiol 2016;30:dyw089. https://doi.org/10.1093/ije/dyw089.

[40] Paillard-Borg S, Wang H-X, Winblad B, Fratiglioni L. Pattern of participation in leisure activities among older people in relation to their health conditions and contextual factors: a survey in a Swedish urban area. Ageing Soc 2009;29:803–21. https://doi.org/10.1017/S0144686X08008337.

[41] Fry A, Littlejohns TJ, Sudlow C, Doherty N, Adamska L, Sprosen T, et al. Comparison of Sociodemographic and Health-Related Characteristics of UK Biobank Participants With Those of the General Population. Am J Epidemiol 2017;186:1026–34. https://doi.org/10.1093/aje/kwx246.

