## Supplementary Materials for "Mid-life and late life activities and their relationship with MRI measures of brain structure and functional connectivity in the UK Biobank cohort"

**Figure S1.** A timeline of the study assessments and variables of interest examined in the current analyses.

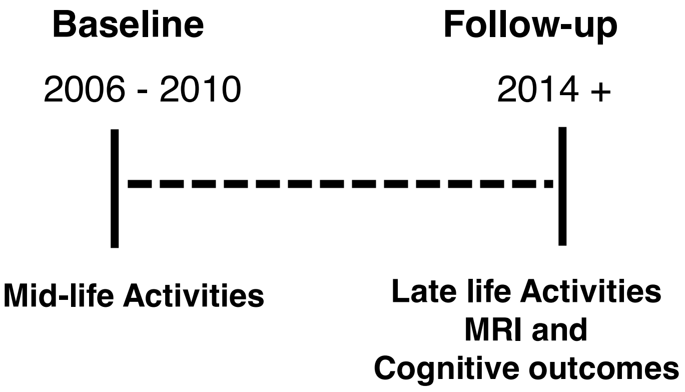


**Abbreviations-** MRI = Magnetic resonance imaging.

**Figure S2.** Flowchart of participant selection and exclusion.

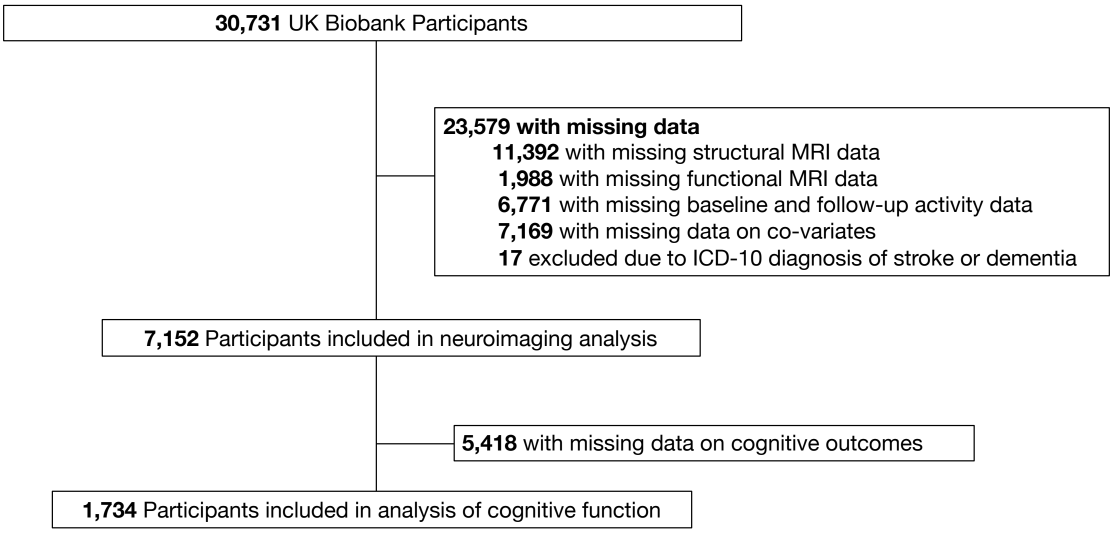

### SUPPLEMENTARY METHODS

#### Activity measures

Respondents were required to indicate from a list of activities, those that were undertaken on a weekly (or more frequent) basis. The activities measured consisted of going to a pub or social club, participating in a religious activity, attending adult education classes and going to a sports club or visiting the gym. A fifth item, which asked individuals to indicate whether they regularly participated in "other group activities" was excluded due to the ambiguity of the question posed. Responses on these four items were coded as a "1" for weekly participation, or a "0" for irregular/ no participation.

Separately, respondents were also asked to indicate how frequently they received or made friend or family visits, with possible answers consisting of: "almost daily", "2-4 times a week", "about once a week", "about once a month", "once every few months" or "never or almost never". Leisure-time computer use was included as another item, with participants asked to indicate the number of hours (per week) they committed to this activity in a typical day. To avoid biases introduced by differences in the granularity of our various activity scales, we binarized responses on these items. Specifically, we assigned a code of "0" for individuals who received or made friend and family visits at least once a week, and an assignment of "0" for those who visited friends and family less frequently than once a week. Weekly or more frequent family/friends were coded as "1". Similarly, participants reporting 0 hours of daily computer use were coded as "0", whereas, those reporting any amount of daily use where grouped as weekly engagers (i.e. “1”). For all activities, answers corresponding to “do not know” or “prefer not to answer” were coded as missing data.

**Cognitive measures**

A cognitive battery designed specifically for UK Biobank was administered via a tablet. While the self-administered and unsupervised nature of these tests may call into question the reliability and validity of these tests, a recent study of 160 adults [1] shows that the UK Biobank cognitive assessments have moderate-to-high test-retest reliability and moderate-to-high concurrent validity, as indicated by correlations between each of these measures with that of validated and standardized tests, delivered in a face-to-face setting. The selection of cognitive tests examined in the present study was guided by prior research [2,3]. Some tests, for instance of language fluency, were only administered at the pilot phase of the UK Biobank, and so were excluded here. Overall, several previous studies have demonstrated that the cognitive tests examined here are sensitive to age-related differences in performance [1,3].

The cognitive battery included a 13-item questionnaire designed to assess Fluid Intelligence. This questionnaire included reasoning/logic items that each had 5 multiple choice options. Participants were required to select one of these responses within a 2-minute period. For example, one of the items asked: “Relaxed means the opposite of?”, with the response options consisting of “calm”, “anxious”, “cool”, “worried” and “tense”. The outcome of this test was the total number of questions answered correctly.

Numeric and alphanumeric trail making tasks were also completed by a portion of the total sample. This task presented participants with 25 circles distributed across the screen, which contained either numbers ranging from 1-25 (numeric trail) or a mix of numbers and letters (alphanumeric trail). Participants were asked to connect the circles from the smallest to largest numbers in one task (numeric trail) or to alternate between numbers and letters in an ascending sequence in the other (alphanumeric trail). The outcomes on these tasks were the time taken to complete the trail, with lower values reflecting a faster completion time. Individuals who spent ≥ 250 seconds on these respective tests, were excluded [4].

A digit span test was also administered. Individuals were presented with a sequence of digits, which they were instructed to recall in a reverse order (starting with the last digit first) after a short delay. Each time the sequence of digits was recalled correctly, the number of digits to be remembered increased by 1. The number began at 2 digits and increased to a maximum number of 12 digits. The test terminated as soon as an error on a given trial was made. The outcome here was the maximum number of digits recalled.

Another test, known as pairs matching, relied on the use of computerized cards. Here, participants were presented with a set of cards (organized into 3 rows, 4 columns) and were briefly shown the symbols that the cards contained before they were turned face down. Each card had a matching pair, with the task requiring respondents to match the six pairs of cards from memory while making as few errors as possible.  The main outcome was the number of incorrect matches made.

The prospective memory test began with a screen containing four coloured symbols: a blue square, a pink star, a grey cross, and an orange circle. Text also accompanied the four symbols that informed participants that they would see the screen again at the end of the cognitive battery, and they would be asked to press the blue square. The text instructed participants that rather than touching the blue square (as the later instructions dictated), they should instead press the orange circle. The outcome was whether or not participants had responded corrected (i.e. touched the orange circle) on the first attempt.

A symbol digit matching task was also administered. In this task, individuals were presented with a set of grids with each box containing a symbol. Participants were required to match each symbol with a unique digit, with the correct symbol-digit matches detailed in a key that was available throughout the 1-minute response period. The outcome was the number of correct symbol-digit matches made (excluding the first 8 items that served as training material).

Finally, a simple reaction time task was also completed, which is analogous to the card game ‘Snap’. For this assessment, participants were presented with a pair of cards (total number of pairs = 12) and were required to press a button as quickly as possible whenever the symbols on the cards were matching. Four training trials were first completed, after which 7 recorded trials were administered (4 of which contained matching pairs). The key outcome was the mean response time (in seconds) across the 4 trials containing matching pairs.

**Demographics and health-related variables**

Educational level was assessed by asking respondents whether they possessed one or more of the following qualifications: “college or university degree” (= “5”), “NVQ or HND or HNC or equivalent” (= “4”), “other professional qualifications (e.g. nursing, teaching)” (= “4”), “A levels/AS levels or equivalent” (= “3”), “O levels/ GCSEs or equivalent” (= “2”) and “CSE or equivalent” (= “1”). From this information, we created a variable indicating the highest qualification earned. Occupational status was coded according to the Standard Occupational Classification, where participants were divided into one of nine occupational groups: “Manager and Senior Officials” (= “9”), “Professional Occupations” (= “8”), “Associate Professional and Technical Occupations” (= “7”), “Administrative and Secretarial Occupations” (= “6”), “Skilled Trades Occupations” (= “5”), “Personal Service Occupations” (= “4”), “Sales and Customer Service Occupations” (= “3”), “Process, plant and machine operatives” (= “2”) and “Elementary Occupations” (= “1”). An additional category, reflecting the lowest level of this variable included those who were retired, unemployed, looking after home and/ or family, unable to work because of sickness or disability, full or part time student (= "0").

Frequency of alcohol intake (over the last year) was measured as "daily or almost daily" (= "5"), "three or four times a week" (= "4"), "once or twice a week" (= "3"), "one to three times a month" (= "2"), "special occasions only" (= "1") or "never" (= "0"). Sleep duration was measured as the hours of sleep within an average 24-hour period. Body mass index (BMI) was calculated as: (weight [kilograms]/height [metres])$^2$. Mean Arterial Pressure (MAP) was also calculated based on the systolic and diastolic indices measured with an electronic tool (systolic blood pressure + 2* diastolic blood pressure/3). Note that in the case where two measurements of diastolic/systolic measures were available, an average over these measures were created before calculating the MAP. If only a single assessment was available, then this was used for to calculate the MAP. Diagnoses of depressive (e.g. major depression) and anxiety disorders (e.g. social anxiety), coded according to the World Health Organization's International Classification of Diseases manual (ICD-10), over the study period were also taken into account. Until April 2010, ICD-10 was used, with ICD-10 4th edition used from April 2010 to date [5]. An index of social isolation was further included [6], assessed as the total number of individuals living in the household (alongside the participant).

**MRI Data Acquisition**

T1 images were acquired with a resolution of 1 x 1 x 1 mm and a field of view of 256 mm, in the sagittal plane with a 3D magnetization-prepared rapid gradient echo (MPRAGE). The inversion and repetition times were 880 ms and 2000 ms, respectively. Diffusion-weighted images were collected using a spin-echo echo planar imaging sequence, with 2 mm isotropic voxels, a 104 x 104 mm field of view, an echo time of 92 ms, repetition time of 3600 ms and a multiband acceleration factor of 3 (i.e. three slices acquired at a time [7]. Five b0 images were collected, in addition to images acquired with two separate b-values (50x b = 1000 s/mm$^2$ and 50x b = 2,0000 s/mm$^2$), which amounted to 100 diffusion-encoding directions. T2-weighted FLAIR imaging was additionally acquired with 3D SPACE in the sagittal plane (resolution = 1.05 x 1 x 1 mm, field of view = 192 x 256 x 256 mm; inversion time = 1800 ms, repetition time = 5000 ms). Finally, resting-state fMRI images were acquired with a gradient echo-echo planar imaging (GE-EPI) using a multi-slice acceleration of 8 and a flip angle of 52° (2.4 x 2.4 x 2.4 mm voxels; field of view = 88 x 88 x 64 mm, 490 timepoints, repetition time = 0.745 seconds, echo time = 39 ms).

**MRI Data Pre-processing**

Measures of brain structure and functional connectivity, or Image Derived Phenotypes (IDPs), were generated using FMRIB’s Biobank Pipeline (version 1.0, [8]). For a detailed description of the imaging protocol and pre-processing steps, please see Alfaro-Almagro et al. [8] and Smith et al.[9].

The T1-weighted images were first defaced to anonymize the images. The size of field of view was then reduced to remove voxels containing non-brain tissue, using a combination of FSL's Brain Extraction Tool [10]; linear registration [11,12], and the MNI152 "nonlinear 6$^{th}$ generation" standard space T1 template (http://www.bic.mni.mcgill.ca/ServicesAtlases/ICBM152NLin6). Gradient distortion correction was also applied at this point. Next, nonlinear registration was performed with FNIRT [13] to calculate the T1-to-MNI152 warp transform, with a custom brain mask as a reference image. Using the inverse of the warp transform, the standard space brain mask was then transformed into each individual's native T1 space in order to brain extract for each individual. FAST [14] was subsequently applied to fulfil two objectives on the brain extracted T1 images: (1) segment the images based on tissue types (i.e. GM, WM and CSF) and (2) generate a bias-field corrected image. Partial volume estimates of GM from FAST were parcellated into 139 GM ROIs, using a combination of the Harvard-Oxford cortical and subcortical atlases (https://fsl.fmrib.ox.ac.uk/fsl/fslwiki/Atlases) and Diedrichsen cerebellar atlas (http://www.diedrichsenlab.org/imaging/propatlas.htm) to facilitate parcellation. FSL's FIRST was also employed to extract volumetric estimates of key subcortical structures, including the hippocampus, amygdala, thalamus, pallidum, caudate, putamen [15]. Note that the corresponding FAST-extracted ROIs for these subcortical regions were excluded. For a list of all of the regional GM IDPs examined in this study, see Appendix 5.2. Finally, in order to derive an estimate of head size (used as a co-variate in the present analyses), the pre-processed T1 images were separately submitted to a

SIENAX (Structural Image Evaluation, using Normalisation, of Atrophy: Cross-sectional; [16].

For the raw diffusion-weighted images, EPI distortions and eddy currents/outlier slices were addressed with FSI's topup [17] and eddy, respectively [18,19]. The next step was to apply gradient distortion correction (developed by HCP and FSL) to remove artefacts introduced by head motion. DTI fit [20] was applied to derive FA and MD images for each participant. The pre-processed diffusion-weighted images were subsequently submitted to a tractography-based analysis. Specifically, BEDPOSTX (Bayesian Estimation of Diffusion Parameters Obtained using Sampling Techniques, http://fsl.fmrib.ox.ac.uk/fsl/fslwiki/FDT/UserGuide) was used, which is a method for intra-voxel modelling of multi-fibre tract orientations with the capacity to estimate up to three fibre orientations within a voxel. The output of BEDPOSTX were then fed to PROBTRACKX [21–24], a tool that conducts probabilistic tractography (implemented with a crossing-fibre model), which is currently able to map 27 major tracts using start/stop ROI masks defined by AutoPX [25]. The tracts included the cingulum bundle, thalamic radiations, longitudinal fasciculi with a full list available in the Appendix 5.3. As the results of both of these tools are in native space, non-linear transformations were applied to bring the results into 1mm standard MNI space. The IDPs made available after these pre-processing steps were the weighted-mean FA and MD for each of the 27 tracts. After defacing, the T2 FLAIR structural images were registered to the T1-weighted images using FLIRT [26]. The transforms generated were then used to register the FLAIR images to MNI space. An estimate of total WM hyperintensity volume was generated by feeding the T2-weighted FLAIR and T1-weighted images to the BIANCA tool developed by Griffanti et al. [27].

The pre-processing steps involved motion correction (via MCFLIRT, [12]), grand-mean intensity normalisation of the 3D dataset, high pass temporal filtering, EPI and GDC unwarping. ICA+FIX processing were applied to remove the presence of individual-level structured artefacts [28–30]. The images were then brought into T1 space and then standard MNI space, using FLIRT (with a BBR cost function [26]. Next, low-dimensionality Group ICA (n. of components = 25) was applied to the pre-processed functional images of 4,162 individuals [8] in order to derive group-level spatial maps of large-scale resting state networks. Twenty-one of these components were deemed to be 'signal' of interest (see Figure S3 for an overview of components; hand labelled by Melis Anatürk (MA), Sana Suri (SS) and Claire Sexton (CES) with disagreements resolved through discussion). These components were then submitted to a FSLNETS analysis. This tool was used to generate a 21x21 matrix (i.e. connectomes) representing the correlations (i.e. edge or connectivity) between pairs of large-scale functional networks (i.e. nodes). Partial correlations were used as they provide a more direct estimate of the connectivity between two nodes, with estimates adjusted for the connectivity of all other nodes in the connectome. Estimates of partial correlations were derived with an L2 Regularization (rho = 0.05 in the Ridge Regression option in FSL Nets). All values were transformed from Pearson's correlations to z-statistics. Prior to the analysis, we used a method validated by prior studies [31,32] to improve the interpretability of our results. Specifically, values for individuals were multiplied by the sign of their mean edge value, in order to give an index of *absolute connectivity,* with higher values reflecting stronger connections. Head motion (derived from McFLIRT) represented the mean relative displacement acquired during the acquisition of functional images and was used here as a co-variate.

**FDR Corrections**

FDR corrections were applied across imaging modalities, for activities measured at the same time-point (mid-life; late life). FDR corrections were applied separately for the analysis of cognitive tests, as this was conducted in a smaller sample to the main analytical group.

**Figure S3. Classifications of signal components from a 25-dimension ICA.**

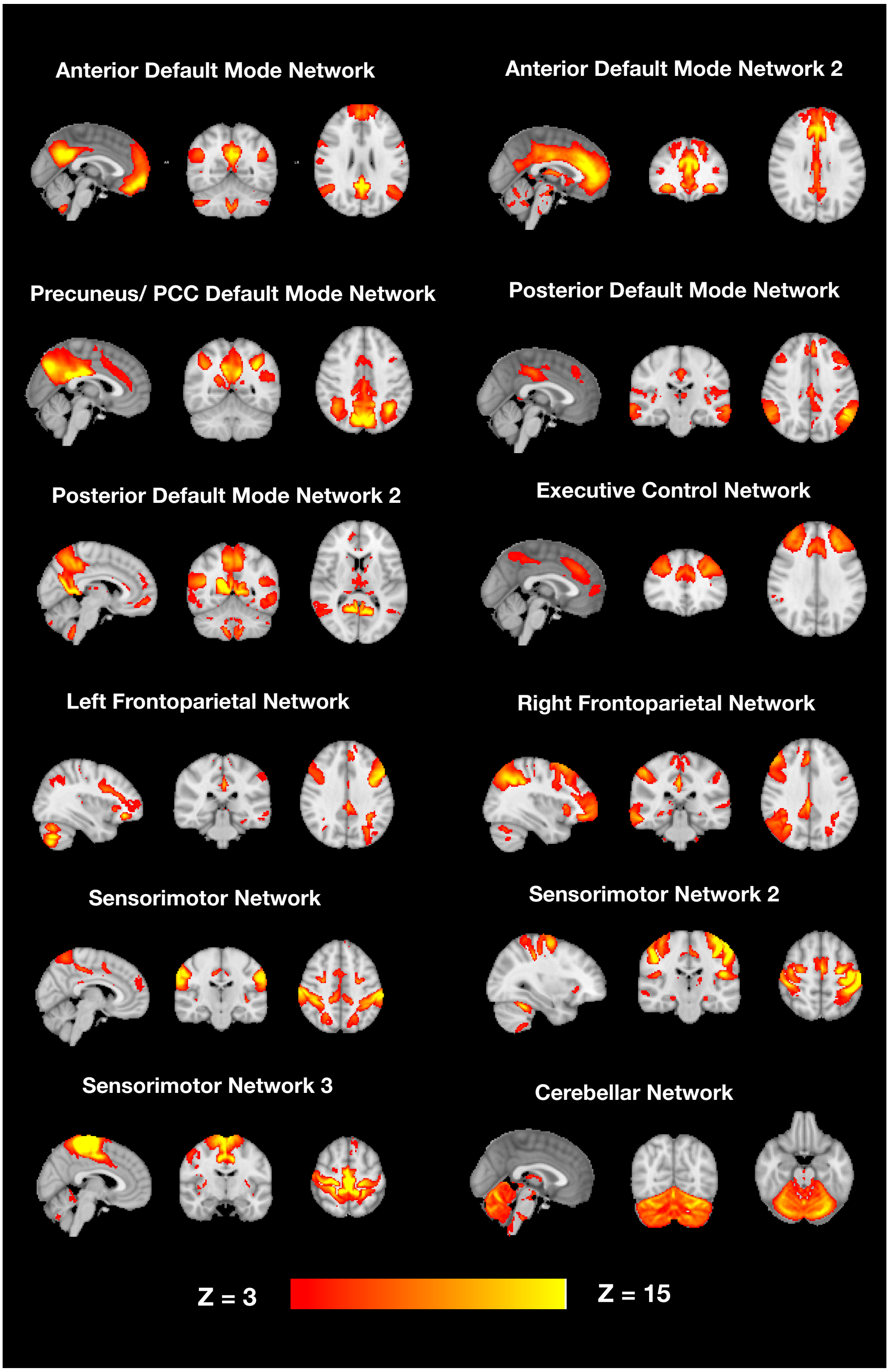


**Abbreviations** – PCC = Posterior Cingulate Cortex.

**Figure S3 continued. Classifications of signal components from a 25-dimension ICA.**

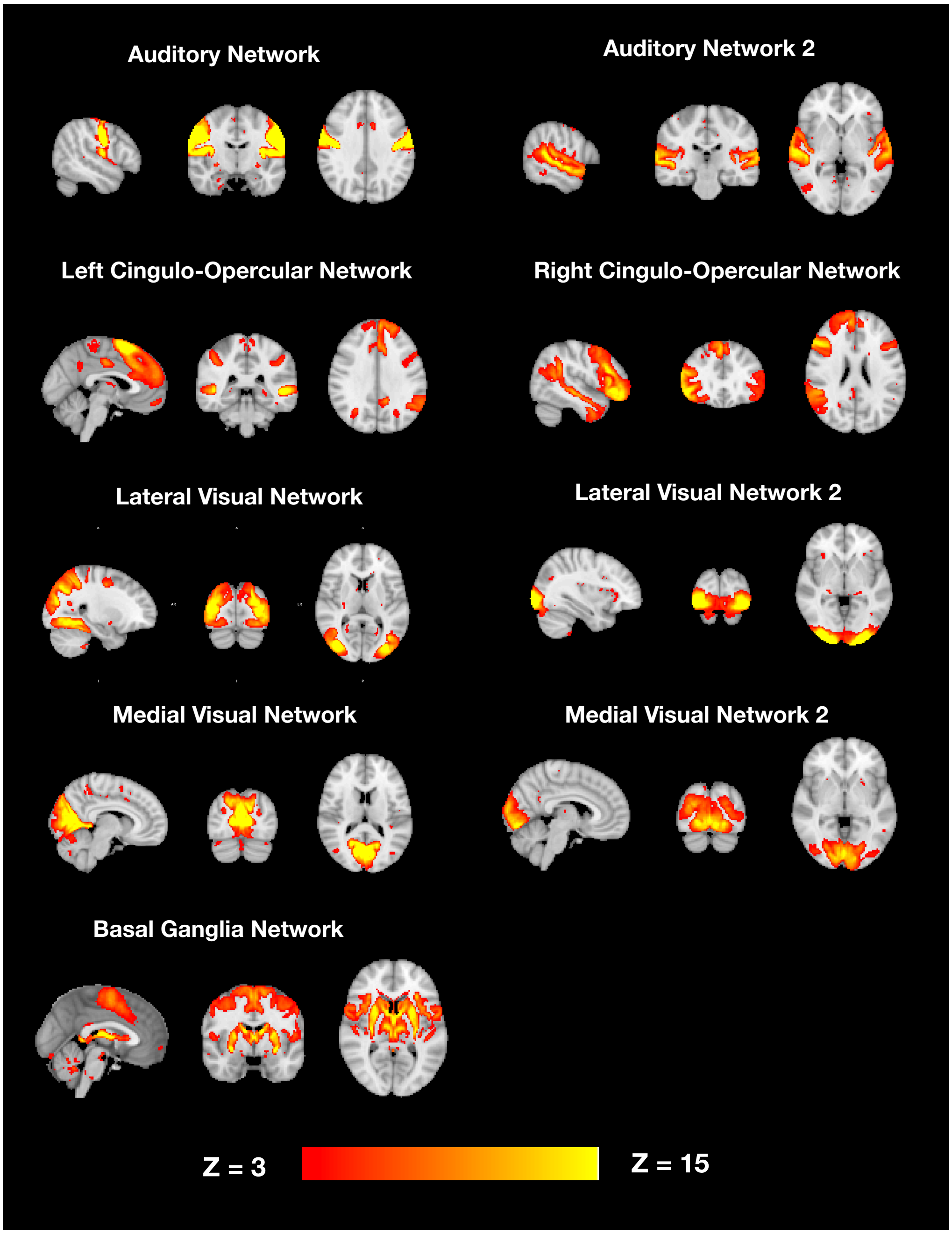

### SUPPLEMENTARY RESULTS

#### Comparisons of included and excluded individuals

For a comparison of included and excluded participants, see Table S4. Included participants were on average, older, more likely to be female, more educated, from a higher occupational grade and had a lower BMI (all FDR q-values < 0.01). They also consumed, on average, more alcohol, lived with fewer people and slept longer hours per night (all FDR q-values < 0.01). Included individuals had less relative head motion (FDR q-value < 0.01) than those excluded. There were no differences in BP, head size or the number of depressed/anxious individuals (FDR q-value > 0.01).

**Table S1. Comparisons between included and excluded individuals.**

| Dependent variable | Included | Excluded | Test-statistic | p-value | FDR q-value |
|---|---|---|---|---|---|
| Age at baseline (years) | 54.571 | 56.385 | t = -18.286 | < 0.001 | < 0.001 |
| Sex (% female) | 4897 (54.48%) | 4897 (51.05%) | χ2 = 25.925 | < 0.001 | < 0.001 |
| Educational level | 4.071 | 4.174 | t = -6.914 | < 0.001 | < 0.001 |
| Occupational status | 4.702 | 6.105 | t = -29.068 | < 0.001 | < 0.001 |
| Alcohol (frequency/ week) | 3.257 | 3.414 | t = -8.541 | < 0.001 | < 0.001 |
| BMI ($kg/m^2$) | 26.821 | 26.4 | t = 7.577 | < 0.001 | < 0.001 |
| N. in household | 2.556 | 2.442 | t = 7.433 | < 0.001 | < 0.001 |
| Sleep duration (hours/ night) | 7.148 | 7.215 | t = -5.284 | < 0.001 | < 0.001 |
| Head motion | 0.123 | 0.119 | t = 4.965 | < 0.001 | < 0.001 |
| % of individuals with ICD-10 diagnosis of Depression/ Anxiety | 51 (0.71%) | 181 (0.78%) | χ2 = 0.218 | 0.641 | 0.76 |
| BP (MAP) | 99.512 | 99.575 | t = -0.39 | 0.697 | 0.76 |
| Head size | 1.299 | 1.299 | t = 0.144 | 0.886 | 0.89 |

**Abbreviations-** BMI = Body mass index; BP = Blood pressure; ICD = International Classification of Diseases; MAP = Mean arterial pressure; N = number.
**Note:** Due to unequal variances detected between included and excluded participants on continuous variables, Welch's t-test was performed and is reported.

**Table S2. Table of associations between weekly mid-life participation in activities and cognitive function.** Results are adjusted for sociodemographic, health and lifestyle co-variates.

| Dependent variable | Predictor | B | SE | B | p-value | FDR q-value |
|---|---|---|---|---|---|---|
| **Numeric Trail making (sec, log)** | **Mid-life computer use** | **-0.040** | **0.009** | **-0.105** | **< 0.001** | **< 0.001** |
| **Alphanumeric trail making (sec, log)** | **Mid-life computer use** | **-0.032** | **0.010** | **-0.068** | **0.002** | **0.036** |
| Fluid intelligence score | Mid-life attendance to social clubs or pubs | -0.226 | 0.102 | -0.056 | 0.027 | 0.299 |
| Simple reaction time (sec) | Mid-life participation in sports clubs or gyms | -0.010 | 0.005 | -0.049 | 0.037 | 0.299 |
| Fluid intelligence score | Mid-life computer use | 0.288 | 0.146 | 0.046 | 0.049 | 0.299 |
| | | **OR** | **95% C.I.** | | **p-value** | **FDR q-value** |
| **Prospective memory score** | **Mid-life computer use** | **2.009** | **1.336 - 22.971** | | **0.001** | **0.014** |

**Abbreviations-** C.I. = Confidence intervals; OR = Odds ratio.
**Bold = survived FDR corrections.**

**Table S3. Associations between weekly late-life participation in activities and cognitive function.** Results are adjusted for sociodemographic, health and lifestyle co-variates.

| Dependent variable | Predictor | B | SE | B | p-value | FDR q-value |
|---|---|---|---|---|---|---|
| **Fluid intelligence score** | **Late-life computer use** | **0.701** | **0.200** | **0.082** | **< 0.001** | **0.011** |
| **Alphanumeric trail making (sec, log)** | **Late-life computer use** | **-0.049** | **0.014** | **-0.075** | **0.001** | **0.011** |
| **Numeric Trail making (sec, log)** | **Late-life computer use** | **-0.039** | **0.012** | **-0.073** | **0.001** | **0.015** |
| **Fluid intelligence score** | **Late-life attendance to adult educational classes** | **0.463** | **0.146** | **0.075** | **0.002** | **0.015** |
| Fluid intelligence score | Late-life attendance to a social club or pub | -0.259 | 0.102 | -0.064 | 0.011 | 0.087 |
| Alphanumeric trail making (sec, log) | Late-life attendance to adult educational classes | -0.025 | 0.010 | -0.052 | 0.019 | 0.128 |
| Fluid intelligence score | Late-life family and friend visits | -0.246 | 0.116 | -0.049 | 0.034 | 0.205 |
| Backward digit span | Late-life computer use | 0.268 | 0.130 | 0.049 | 0.039 | 0.206 |
| | | **OR** | **95% C.I.** | | **p-value** | **FDR q-value** |
| **Prospective memory score** | **Late-life computer use** | **2.554** | **1.507 - 4.208** | | **> 0.001** | **0.011** |

**Abbreviations-** C.I. = Confidence intervals; OR = Odds ratio.
**Bold = survived FDR corrections.**

**Table S4. Table of associations between weekly mid-life participation in activities and volumetric brain measures.** Results are adjusted for sociodemographic, health and lifestyle co-variates.

| Dependent Variable | Predictor | B | SE | β | p-value | FDR q-value |
|---|---|---|---|---|---|---|
| **volume of putamen (left)** | **Mid-life computer use** | **77.429** | **17.534** | **0.042** | **< 0.001** | **0.012** |
| volume of GM in occipital pole (left) | Mid-life family and friend visits | 141.553 | 36.052 | 0.040 | < 0.001 | 0.053 |
| volume of putamen (right) | Mid-life computer use | 61.071 | 16.999 | 0.033 | < 0.001 | 0.143 |
| volume of GM in v cerebellum (right) | Mid-life family and friend visits | -34.862 | 10.026 | -0.037 | 0.001 | 0.143 |
| volume of GM in paracingulate gyrus (right) | Mid-life family and friend visits | 71.750 | 21.099 | 0.034 | 0.001 | 0.150 |
| volume of GM in vermis viib cerebellum | Mid-life attendance to social clubs or pubs | -2.400 | 0.728 | -0.041 | 0.001 | 0.172 |
| volume of GM in precuneous cortex (right) | Mid-life participation in religious activities | -109.239 | 33.578 | -0.031 | 0.001 | 0.172 |
| volume of accumbens (left) | Mid-life computer use | 13.055 | 4.097 | 0.034 | 0.001 | 0.184 |
| volume of putamen (left) | Mid-life participation in religious activities | -42.560 | 13.709 | -0.031 | 0.002 | 0.197 |
| volume of GM in crus ii cerebellum (right) | Mid-life participation in religious activities | -95.910 | 31.111 | -0.035 | 0.002 | 0.197 |
| volume of GM in occipital pole (right) | Mid-life family and friend visits | 104.911 | 34.452 | 0.031 | 0.002 | 0.211 |
| volume of GM in iiv cerebellum (right) | Mid-life family and friend visits | -25.174 | 8.437 | -0.033 | 0.003 | 0.225 |
| volume of hippocampus (left) | Mid-life computer use | 48.577 | 16.395 | 0.032 | 0.003 | 0.233 |
| volume of GM in ventral striatum (right) | Mid-life computer use | 10.629 | 3.662 | 0.030 | 0.004 | 0.254 |
| volume of GM in inferior frontal gyrus pars triangularis (right) | Mid-life computer use | 50.651 | 17.764 | 0.032 | 0.004 | 0.281 |
| volume of GM in temporal pole (left) | Mid-life computer use | 109.799 | 39.281 | 0.028 | 0.005 | 0.305 |
| volume of GM in occipital fusiform gyrus (right) | Mid-life participation in religious activities | -42.894 | 15.546 | -0.031 | 0.006 | 0.313 |
| volume of GM in middle temporal gyrus anterior division (left) | Mid-life computer use | 35.017 | 12.991 | 0.029 | 0.007 | 0.313 |
| volume of GM in temporal fusiform cortex anterior division (left) | Mid-life computer use | 21.712 | 8.084 | 0.028 | 0.007 | 0.313 |
| volume of GM in ventral striatum (left) | Mid-life computer use | 9.827 | 3.662 | 0.027 | 0.007 | 0.313 |
| volume of caudate (right) | Mid-life attendance to adult educational classes | 35.554 | 13.473 | 0.027 | 0.008 | 0.339 |
| volume of GM in brainstem | Mid-life attendance to adult educational classes | -72.939 | 27.954 | -0.028 | 0.009 | 0.343 |
| volume of GM in temporal pole (right) | Mid-life computer use | 99.217 | 38.061 | 0.026 | 0.009 | 0.343 |
| volume of GM in v cerebellum (left) | Mid-life family and friend visits | -26.550 | 10.207 | -0.028 | 0.009 | 0.343 |
| volume of GM in temporal fusiform cortex anterior division (right) | Mid-life computer use | 19.237 | 7.408 | 0.027 | 0.009 | 0.343 |
| volume of GM in vermis viiia cerebellum | Mid-life participation in religious activities | -12.242 | 4.730 | -0.031 | 0.010 | 0.343 |
| volume of GM in superior temporal gyrus anterior division (left) | Mid-life computer use | 25.851 | 10.024 | 0.028 | 0.010 | 0.343 |
| volume of GM in vermis vi cerebellum | Mid-life attendance to social clubs or pubs | -15.576 | 6.050 | -0.032 | 0.010 | 0.343 |
| volume of GM in vermis crus ii cerebellum | Mid-life participation in sports clubs or gyms | -4.633 | 1.804 | -0.031 | 0.010 | 0.343 |
| volume of GM in brainstem | Mid-life attendance to social clubs or pubs | -53.706 | 20.983 | -0.030 | 0.011 | 0.343 |
| volume of GM in central opercular cortex (left) | Mid-life attendance to adult educational classes | 42.683 | 16.729 | 0.025 | 0.011 | 0.343 |
| volume of GM in parahippocampal gyrus anterior division (left) | Mid-life computer use | 34.155 | 13.489 | 0.025 | 0.011 | 0.343 |

| | | | | | | |
|---|---|---|---|---|---|---|
| volume of GM in supramarginal gyrus anterior division (left) | Mid-life family and friend visits | 45.519 | 17.984 | 0.028 | 0.011 | 0.343 |
| volume of GM in lateral occipital cortex superior division (left) | Mid-life family and friend visits | 141.385 | 55.987 | 0.025 | 0.012 | 0.343 |
| volume of GM in crus ii cerebellum (right) | Mid-life attendance to social clubs or pubs | -72.719 | 28.820 | -0.030 | 0.012 | 0.343 |
| volume of GM in iiv cerebellum (left) | Mid-life family and friend visits | -19.658 | 7.812 | -0.028 | 0.012 | 0.345 |
| volume of GM in crus ii cerebellum (left) | Mid-life participation in religious activities | -78.007 | 31.731 | -0.028 | 0.014 | 0.372 |
| volume of GM in insular cortex (left) | Mid-life computer use | 45.262 | 18.420 | 0.021 | 0.014 | 0.372 |
| volume of GM in insular cortex (right) | Mid-life participation in religious activities | -34.842 | 14.322 | -0.022 | 0.015 | 0.375 |
| volume of caudate (right) | Mid-life participation in sports clubs or gyms | 21.795 | 9.032 | 0.025 | 0.016 | 0.381 |
| volume of GM in temporal fusiform cortex anterior division (left) | Mid-life participation in religious activities | 15.245 | 6.321 | 0.026 | 0.016 | 0.381 |
| volume of GM in vermis crus ii cerebellum | Mid-life attendance to social clubs or pubs | -4.836 | 2.020 | -0.031 | 0.017 | 0.392 |
| volume of putamen (right) | Mid-life participation in religious activities | -31.768 | 13.291 | -0.023 | 0.017 | 0.392 |
| volume of GM in middle temporal gyrus anterior division (right) | Mid-life computer use | 25.226 | 10.614 | 0.026 | 0.017 | 0.392 |
| volume of GM in vermis crus i cerebellum | Mid-life computer use | 0.136 | 0.058 | 0.028 | 0.020 | 0.409 |
| volume of GM in superior temporal gyrus anterior division (right) | Mid-life attendance to adult educational classes | 22.651 | 9.806 | 0.025 | 0.021 | 0.409 |
| volume of GM | Mid-life attendance to social clubs or pubs | -1675.634 | 725.392 | -0.015 | 0.021 | 0.409 |
| volume of caudate (left) | Mid-life attendance to social clubs or pubs | 21.876 | 9.478 | 0.025 | 0.021 | 0.409 |
| volume of GM in insular cortex (right) | Mid-life computer use | 42.193 | 18.318 | 0.020 | 0.021 | 0.409 |
| volume of caudate (left) | Mid-life attendance to adult educational classes | 29.073 | 12.628 | 0.023 | 0.021 | 0.409 |
| volume of GM in vermis viiia cerebellum | Mid-life attendance to social clubs or pubs | -9.976 | 4.382 | -0.028 | 0.023 | 0.421 |
| volume of GM in vi cerebellum (right) | Mid-life family and friend visits | -60.922 | 26.766 | -0.023 | 0.023 | 0.421 |
| volume of GM in precentral gyrus (right) | Mid-life attendance to adult educational classes | 117.008 | 51.424 | 0.023 | 0.023 | 0.421 |
| volume of GM in angular gyrus (right) | Mid-life participation in sports clubs or gyms | -56.025 | 24.736 | -0.025 | 0.024 | 0.427 |
| volume of GM in postcentral gyrus (right) | Mid-life family and friend visits | 84.104 | 37.437 | 0.023 | 0.025 | 0.427 |
| volume of GM in vermis viib cerebellum | Mid-life participation in religious activities | -1.755 | 0.786 | -0.027 | 0.026 | 0.427 |
| volume of GM in frontal operculum cortex (left) | Mid-life family and friend visits | 15.245 | 6.848 | 0.024 | 0.026 | 0.427 |
| volume of GM in ventral striatum (right) | Mid-life participation in sports clubs or gyms | 5.273 | 2.369 | 0.023 | 0.026 | 0.427 |
| volume of GM in parahippocampal gyrus anterior division (right) | Mid-life attendance to adult educational classes | 30.516 | 13.733 | 0.022 | 0.026 | 0.427 |
| volume of amygdala (left) | Mid-life participation in sports clubs or gyms | 12.444 | 5.607 | 0.025 | 0.026 | 0.427 |
| volume of GM in inferior temporal gyrus temporooccipital part (right) | Mid-life participation in religious activities | -43.080 | 19.445 | -0.024 | 0.027 | 0.427 |
| volume of GM in postcentral gyrus (left) | Mid-life family and friend visits | 84.362 | 38.109 | 0.023 | 0.027 | 0.427 |
| volume of GM in planum polare (right) | Mid-life attendance to adult educational classes | 14.431 | 6.525 | 0.022 | 0.027 | 0.427 |
| volume of GM in x cerebellum (left) | Mid-life computer use | 5.976 | 2.704 | 0.024 | 0.027 | 0.427 |
| volume of accumbens (right) | Mid-life computer use | 8.250 | 3.745 | 0.024 | 0.028 | 0.427 |
| volume of GM in frontal pole (left) | Mid-life family and friend visits | 126.062 | 57.371 | 0.018 | 0.028 | 0.428 |
| volume of GM in lingual gyrus (right) | Mid-life attendance to adult educational classes | 54.488 | 25.123 | 0.020 | 0.030 | 0.443 |

| | | | | | | |
|---|---|---|---|---|---|---|
| volume of GM in vermis crus i cerebellum | Mid-life participation in religious activities | -0.097 | 0.046 | -0.027 | 0.033 | 0.461 |
| volume of hippocampus (left) | Mid-life participation in religious activities | -27.208 | 12.819 | -0.024 | 0.034 | 0.461 |
| volume of GM in cingulate gyrus anterior division (left) | Mid-life attendance to adult educational classes | 72.624 | 34.527 | 0.023 | 0.035 | 0.461 |
| volume of GM in middle temporal gyrus anterior division (left) | Mid-life participation in religious activities | 21.342 | 10.157 | 0.023 | 0.036 | 0.461 |
| volume of GM in crus i cerebellum (right) | Mid-life attendance to social clubs or pubs | -87.569 | 41.697 | -0.024 | 0.036 | 0.461 |
| volume of GM in temporal fusiform cortex posterior division (left) | Mid-life computer use | 36.005 | 17.147 | 0.020 | 0.036 | 0.461 |
| volume of GM in vermis viiia cerebellum | Mid-life family and friend visits | -10.050 | 4.815 | -0.023 | 0.037 | 0.469 |
| volume of GM in middle temporal gyrus anterior division (left) | Mid-life participation in sports clubs or gyms | 17.511 | 8.403 | 0.023 | 0.037 | 0.469 |
| volume of GM | Mid-life attendance to adult educational classes | 2009.258 | 966.394 | 0.012 | 0.038 | 0.469 |
| volume of GM in superior temporal gyrus posterior division (left) | Mid-life attendance to adult educational classes | 32.485 | 15.800 | 0.022 | 0.040 | 0.469 |
| volume of GM in superior temporal gyrus anterior division (left) | Mid-life participation in religious activities | 16.098 | 7.838 | 0.023 | 0.040 | 0.469 |
| volume of GM in viiia cerebellum (left) | Mid-life attendance to social clubs or pubs | -34.926 | 17.023 | -0.024 | 0.040 | 0.469 |
| volume of GM in inferior temporal gyrus temporooccipital part (left) | Mid-life participation in sports clubs or gyms | -27.559 | 13.434 | -0.023 | 0.040 | 0.469 |
| volume of GM in frontal orbital cortex (right) | Mid-life participation in sports clubs or gyms | 29.574 | 14.421 | 0.020 | 0.040 | 0.469 |
| volume of GM in crus i cerebellum (right) | Mid-life participation in religious activities | -92.171 | 45.012 | -0.023 | 0.041 | 0.469 |
| volume of GM in vi cerebellum (left) | Mid-life family and friend visits | -56.654 | 27.728 | -0.021 | 0.041 | 0.469 |
| volume of GM | Mid-life participation in religious activities | -1596.327 | 783.057 | -0.013 | 0.042 | 0.469 |
| volume of GM in parahippocampal gyrus posterior division (right) | Mid-life attendance to adult educational classes | 11.883 | 5.842 | 0.022 | 0.042 | 0.469 |
| volume of hippocampus (right) | Mid-life computer use | 33.537 | 16.521 | 0.022 | 0.042 | 0.469 |
| volume of GM in paracingulate gyrus (left) | Mid-life family and friend visits | 41.866 | 20.634 | 0.020 | 0.042 | 0.469 |
| volume of GM in x cerebellum (right) | Mid-life computer use | 5.414 | 2.670 | 0.022 | 0.043 | 0.469 |
| volume of thalamus (right) | Mid-life participation in religious activities | -29.767 | 14.700 | -0.018 | 0.043 | 0.469 |
| volume of GM in inferior frontal gyrus pars triangularis (left) | Mid-life family and friend visits | 33.177 | 16.535 | 0.023 | 0.045 | 0.472 |
| volume of GM in frontal operculum cortex (left) | Mid-life participation in sports clubs or gyms | 11.017 | 5.566 | 0.022 | 0.048 | 0.484 |
| volume of GM in crus ii cerebellum (left) | Mid-life attendance to social clubs or pubs | -57.983 | 29.394 | -0.024 | 0.049 | 0.485 |
| volume of GM in central opercular cortex (right) | Mid-life attendance to adult educational classes | 34.439 | 17.497 | 0.019 | 0.049 | 0.487 |
| volume of GM in inferior temporal gyrus anterior division (left) | Mid-life computer use | 21.060 | 10.715 | 0.022 | 0.049 | 0.488 |
| volume of white matter | Mid-life participation in religious activities | -1768.355 | 903.981 | -0.012 | 0.050 | 0.493 |

**Abbreviations-** GM = Grey matter.
**Bold = survived FDR corrections.**

**Table S5. Table of associations between weekly late-life participation in activities and volumetric brain measures.** Results are adjusted for sociodemographic, imaging, health and lifestyle co-variates.

| **Dependent Variable** | **Predictor** | **B** | **SE** | **β** | **p-value** | **FDR q-value** |
|---|---|---|---|---|---|---|
| volume of hippocampus (left) | Late-life computer use | 80.416 | 23.304 | 0.038 | 0.001 | 0.256 |
| volume of putamen (left) | Late-life participation in religious activities | -45.291 | 13.557 | -0.033 | 0.001 | 0.256 |
| volume of thalamus (left) | Late-life participation in sports clubs or gyms | 39.043 | 12.873 | 0.026 | 0.002 | 0.345 |
| volume of putamen (right) | Late-life participation in religious activities | -39.007 | 13.138 | -0.029 | 0.003 | 0.345 |
| volume of GM in precuneous cortex (left) | Late-life computer use | -165.336 | 56.679 | -0.027 | 0.004 | 0.345 |
| volume of GM in lateral occipital cortex inferior division (right) | Late-life attendance to adult educational classes | 124.161 | 43.191 | 0.029 | 0.004 | 0.345 |
| volume of GM in occipital pole (left) | Late-life family and friend visits | 103.865 | 37.253 | 0.029 | 0.005 | 0.345 |
| volume of GM in temporal fusiform cortex posterior division (left) | Late-life computer use | 67.507 | 24.363 | 0.027 | 0.006 | 0.345 |
| volume of GM in inferior temporal gyrus posterior division (right) | Late-life attendance to adult educational classes | -70.249 | 25.450 | -0.029 | 0.006 | 0.345 |
| volume of GM in insular cortex (left) | Late-life computer use | 72.265 | 26.189 | 0.024 | 0.006 | 0.345 |
| volume of hippocampus (right) | Late-life computer use | 64.636 | 23.482 | 0.030 | 0.006 | 0.345 |
| volume of GM in precuneous cortex (right) | Late-life participation in religious activities | -89.811 | 33.178 | -0.026 | 0.007 | 0.357 |
| volume of GM in precentral gyrus (left) | Late-life computer use | 203.124 | 75.920 | 0.026 | 0.007 | 0.357 |
| volume of pallidum (right) | Late-life attendance to social clubs or pubs | -15.734 | 5.884 | -0.031 | 0.008 | 0.357 |
| volume of GM in inferior frontal gyrus pars opercularis (left) | Late-life attendance to social clubs or pubs | -34.813 | 13.400 | -0.031 | 0.009 | 0.357 |
| volume of GM in v cerebellum (right) | Late-life computer use | 46.162 | 17.918 | 0.027 | 0.010 | 0.363 |
| volume of caudate (left) | Late-life computer use | 47.645 | 18.615 | 0.025 | 0.011 | 0.363 |
| volume of GM in planum polare (right) | Late-life participation in sports clubs or gyms | -11.151 | 4.373 | -0.025 | 0.011 | 0.363 |
| volume of GM in occipital pole (right) | Late-life family and friend visits | 89.521 | 35.600 | 0.025 | 0.012 | 0.369 |
| volume of white matter | Late-life family and friend visits | 2365.984 | 950.375 | 0.015 | 0.013 | 0.377 |
| volume of thalamus (right) | Late-life participation in sports clubs or gyms | 29.946 | 12.149 | 0.021 | 0.014 | 0.399 |
| volume of GM in inferior temporal gyrus temporooccipital part (right) | Late-life attendance to adult educational classes | 62.069 | 25.609 | 0.025 | 0.015 | 0.437 |
| volume of GM in frontal operculum cortex (left) | Late-life computer use | 29.463 | 12.236 | 0.026 | 0.016 | 0.447 |
| volume of GM in inferior temporal gyrus temporooccipital part (right) | Late-life attendance to social clubs or pubs | 42.792 | 17.776 | 0.027 | 0.016 | 0.447 |
| volume of GM in occipital fusiform gyrus (right) | Late-life participation in religious activities | -36.687 | 15.361 | -0.027 | 0.017 | 0.460 |
| volume of GM in x cerebellum (right) | Late-life computer use | 8.889 | 3.796 | 0.025 | 0.019 | 0.499 |
| volume of GM in frontal operculum cortex (right) | Late-life computer use | 27.169 | 11.646 | 0.025 | 0.020 | 0.506 |

| | | | | | | |
|---|---|---|---|---|---|---|
| volume of GM in vermis crus i cerebellum | Late-life family and friend visits | 0.111 | 0.048 | 0.028 | 0.020 | 0.510 |
| volume of GM in insular cortex (right) | Late-life computer use | 60.129 | 26.041 | 0.020 | 0.021 | 0.517 |
| volume of GM in middle temporal gyrus anterior division (left) | Late-life participation in religious activities | 22.847 | 10.040 | 0.025 | 0.023 | 0.542 |
| volume of GM in vi cerebellum (right) | Late-life computer use | 107.390 | 47.821 | 0.023 | 0.025 | 0.550 |
| volume of GM in vermis viib cerebellum | Late-life attendance to social clubs or pubs | -1.616 | 0.720 | -0.028 | 0.025 | 0.550 |
| volume of GM in frontal operculum cortex (right) | Late-life participation in sports clubs or gyms | 11.821 | 5.297 | 0.025 | 0.026 | 0.562 |
| volume of GM in ventral striatum (right) | Late-life attendance to social clubs or pubs | -5.789 | 2.621 | -0.024 | 0.027 | 0.564 |
| volume of GM in frontal pole (left) | Late-life attendance to adult educational classes | -160.275 | 74.311 | -0.018 | 0.031 | 0.602 |
| volume of GM in v cerebellum (left) | Late-life computer use | 38.963 | 18.235 | 0.023 | 0.033 | 0.613 |
| volume of GM in planum polare (left) | Late-life participation in sports clubs or gyms | -9.258 | 4.349 | -0.022 | 0.033 | 0.621 |
| volume of GM in iiv cerebellum (right) | Late-life computer use | 31.983 | 15.075 | 0.023 | 0.034 | 0.623 |
| volume of GM in temporal fusiform cortex anterior division (right) | Late-life participation in sports clubs or gyms | -10.052 | 4.792 | -0.022 | 0.036 | 0.624 |
| volume of GM in planum temporale (right) | Late-life participation in sports clubs or gyms | -13.541 | 6.456 | -0.021 | 0.036 | 0.624 |
| volume of GM in vermis vi cerebellum | Late-life computer use | 24.546 | 11.876 | 0.023 | 0.039 | 0.652 |
| volume of GM in superior temporal gyrus posterior division (right) | Late-life family and friend visits | 29.600 | 14.344 | 0.021 | 0.039 | 0.652 |
| volume of GM in insular cortex (right) | Late-life participation in sports clubs or gyms | 24.434 | 11.844 | 0.018 | 0.039 | 0.652 |
| volume of GM in inferior temporal gyrus temporooccipital part (right) | Late-life participation in religious activities | -39.561 | 19.195 | -0.023 | 0.039 | 0.652 |
| volume of GM in ventral striatum (left) | Late-life attendance to social clubs or pubs | -5.364 | 2.621 | -0.022 | 0.041 | 0.665 |
| volume of GM in postcentral gyrus (right) | Late-life family and friend visits | 79.061 | 38.679 | 0.021 | 0.041 | 0.665 |
| volume of GM in temporal pole (left) | Late-life computer use | 114.102 | 55.857 | 0.020 | 0.041 | 0.665 |
| volume of GM in vi cerebellum (left) | Late-life computer use | 100.957 | 49.542 | 0.021 | 0.042 | 0.666 |
| volume of GM in juxtapositional lobule cortex formerly supplementary motor cortex (left) | Late-life attendance to social clubs or pubs | 27.425 | 13.776 | 0.023 | 0.047 | 0.701 |
| volume of GM in x cerebellum (left) | Late-life attendance to adult educational classes | -5.502 | 2.788 | -0.021 | 0.048 | 0.701 |
| volume of GM in vermis crus i cerebellum | Late-life attendance to social clubs or pubs | -0.082 | 0.042 | -0.025 | 0.049 | 0.701 |

**Abbreviations-** GM = Grey matter.
**Bold = survived FDR corrections.**

**Table S6. Table to show associations between weekly mid-life participation in activities and tract-averaged FA and MD.** Results are adjusted for sociodemographic, imaging, health and lifestyle co-variates.

| Dependent Variable | Predictor | B | SE | β | p-value | FDR q-value |
|---|---|---|---|---|---|---|
| FA in anterior thalamic radiation (right) | Mid-life computer use | -0.002 | 0.001 | -0.036 | 0.002 | 0.197 |
| FA in anterior thalamic radiation (left) | Mid-life computer use | -0.002 | 0.001 | -0.034 | 0.002 | 0.217 |
| FA in cingulate gyrus part of cingulum (left) | Mid-life computer use | -0.004 | 0.001 | -0.034 | 0.004 | 0.254 |
| FA in superior thalamic radiation (left) | Mid-life computer use | -0.002 | 0.001 | -0.034 | 0.004 | 0.264 |
| FA in corticospinal (left) | Mid-life computer use | -0.002 | 0.001 | -0.029 | 0.013 | 0.360 |
| FA in superior thalamic radiation (right) | Mid-life computer use | -0.002 | 0.001 | -0.028 | 0.015 | 0.375 |
| MD in superior longitudinal fasciculus (right) | Mid-life participation in sports clubs or gyms | 0.000 | 0.000 | 0.028 | 0.015 | 0.375 |
| FA in superior longitudinal fasciculus (left) | Mid-life computer use | -0.002 | 0.001 | -0.027 | 0.018 | 0.395 |
| FA in acoustic radiation (left) | Mid-life computer use | -0.002 | 0.001 | -0.028 | 0.019 | 0.399 |
| MD in middle cerebellar peduncle | Mid-life computer use | 0.000 | 0.000 | -0.027 | 0.019 | 0.404 |
| MD in parahippocampal part of cingulum (right) | Mid-life family and friend visits | 0.000 | 0.000 | -0.026 | 0.025 | 0.427 |
| MD in acoustic radiation (right) | Mid-life computer use | 0.000 | 0.000 | 0.026 | 0.027 | 0.427 |
| MD in cingulate gyrus part of cingulum (right) | Mid-life attendance to adult educational classes | 0.000 | 0.000 | 0.025 | 0.032 | 0.458 |
| MD in forceps minor | Mid-life participation in religious activities | 0.000 | 0.000 | -0.025 | 0.042 | 0.469 |
| FA in anterior thalamic radiation (right) | Mid-life attendance to adult educational classes | -0.001 | 0.001 | -0.023 | 0.043 | 0.469 |
| FA in cingulate gyrus part of cingulum (right) | Mid-life family and friend visits | 0.002 | 0.001 | 0.023 | 0.046 | 0.475 |
| MD in uncinate fasciculus (left) | Mid-life attendance to social clubs or pubs | 0.000 | 0.000 | 0.025 | 0.047 | 0.481 |
| MD in forceps major | Mid-life attendance to adult educational classes | 0.000 | 0.000 | -0.024 | 0.050 | 0.492 |

**Abbreviations-** FA = Fractional Anisotropy; MD = Medial Diffusivity.
**Bold = survived FDR corrections.**

**Table S7. A table to demonstrate associations between weekly late-life participation in activities and tract-averaged FA and MD.** Results are adjusted for sociodemographic, imaging, health and lifestyle co-variates.

| Dependent Variable | Predictor | B | SE | β | p-value | FDR q-value |
|---|---|---|---|---|---|---|
| FA in anterior thalamic radiation (left) | Late-life attendance to adult educational classes | -0.002 | 0.001 | -0.034 | 0.003 | 0.345 |
| FA in anterior thalamic radiation (right) | Late-life attendance to adult educational classes | -0.002 | 0.001 | -0.034 | 0.003 | 0.345 |
| MD in cingulate gyrus part of cingulum (right) | Late-life attendance to adult educational classes | 0.000 | 0.000 | 0.033 | 0.005 | 0.345 |
| MD in superior longitudinal fasciculus (left) | Late-life attendance to adult educational classes | 0.000 | 0.000 | 0.030 | 0.008 | 0.357 |
| MD in inferior frontooccipital fasciculus (right) | Late-life attendance to adult educational classes | 0.000 | 0.000 | 0.030 | 0.008 | 0.357 |
| MD in superior longitudinal fasciculus (right) | Late-life attendance to adult educational classes | 0.000 | 0.000 | 0.030 | 0.009 | 0.357 |
| MD in cingulate gyrus part of cingulum (left) | Late-life attendance to adult educational classes | 0.000 | 0.000 | 0.031 | 0.009 | 0.357 |
| FA in forceps minor | Late-life attendance to adult educational classes | -0.002 | 0.001 | -0.030 | 0.009 | 0.357 |
| MD in anterior thalamic radiation (left) | Late-life attendance to adult educational classes | 0.000 | 0.000 | 0.027 | 0.010 | 0.362 |
| MD in anterior thalamic radiation (right) | Late-life attendance to adult educational classes | 0.000 | 0.000 | 0.027 | 0.011 | 0.363 |
| MD in middle cerebellar peduncle | Late-life computer use | 0.000 | 0.000 | -0.029 | 0.011 | 0.365 |
| MD in inferior frontooccipital fasciculus (left) | Late-life attendance to adult educational classes | 0.000 | 0.000 | 0.027 | 0.018 | 0.467 |
| FA in middle cerebellar peduncle | Late-life participation in religious activities | -0.002 | 0.001 | -0.028 | 0.022 | 0.535 |
| FA in inferior frontooccipital fasciculus (right) | Late-life attendance to adult educational classes | -0.002 | 0.001 | -0.027 | 0.022 | 0.535 |
| FA in inferior frontooccipital fasciculus (left) | Late-life attendance to adult educational classes | -0.002 | 0.001 | -0.026 | 0.024 | 0.542 |
| MD in inferior longitudinal fasciculus (right) | Late-life attendance to adult educational classes | 0.000 | 0.000 | 0.025 | 0.026 | 0.562 |
| MD in middle cerebellar peduncle | Late-life participation in sports clubs or gyms | 0.000 | 0.000 | -0.026 | 0.026 | 0.562 |
| MD in uncinate fasciculus (left) | Late-life participation in religious activities | 0.000 | 0.000 | -0.026 | 0.028 | 0.568 |
| FA in acoustic radiation (right) | Late-life computer use | -0.002 | 0.001 | -0.025 | 0.032 | 0.602 |
| FA in anterior thalamic radiation (left) | Late-life family and friend visits | 0.001 | 0.001 | 0.024 | 0.032 | 0.610 |
| FA in superior thalamic radiation (left) | Late-life computer use | -0.002 | 0.001 | -0.024 | 0.036 | 0.624 |
| MD in acoustic radiation (left) | Late-life attendance to social clubs or pubs | 0.000 | 0.000 | 0.026 | 0.044 | 0.674 |
| MD in forceps minor | Late-life attendance to adult educational classes | 0.000 | 0.000 | 0.024 | 0.044 | 0.674 |

**Abbreviations-** FA = Fractional Anisotropy; MD = Medial Diffusivity.
**Bold = survived FDR corrections.**

**Table S8. Table to show the associations between weekly mid-life participation in activities and resting-state functional connectivity.** Results are adjusted for sociodemographic, imaging, health and lifestyle co-variates.

| Dependent Variable | Predictor | B | SE | β | p-value | FDR q-value |
|---|---|---|---|---|---|---|
| **Connectivity between SMN 3 and Lateral VN 2** | **Mid-life participation in sports clubs or gyms** | **0.063** | **0.014** | **0.056** | **< 0.001** | **0.009** |
| **Connectivity between SMN 3 and CN** | **Mid-life participation in sports clubs or gyms** | **0.062** | **0.014** | **0.051** | **< 0.001** | **0.015** |
| Connectivity between SMN and Posterior DMN | Mid-life attendance to social clubs or pubs | 0.085 | 0.024 | 0.046 | 0.000 | 0.143 |
| Connectivity between L FPN and AN | Mid-life family and friend visits | 0.055 | 0.016 | 0.041 | 0.001 | 0.143 |
| Connectivity between AN and CN | Mid-life participation in sports clubs or gyms | 0.045 | 0.013 | 0.042 | 0.001 | 0.143 |
| Connectivity between Medial VN and R CON | Mid-life attendance to social clubs or pubs | 0.049 | 0.014 | 0.045 | 0.001 | 0.143 |
| Connectivity between SMN 3 and BN | Mid-life participation in sports clubs or gyms | -0.052 | 0.015 | -0.040 | 0.001 | 0.169 |
| Connectivity between Posterior DMN and SMN 3 | Mid-life participation in sports clubs or gyms | 0.043 | 0.013 | 0.040 | 0.001 | 0.170 |
| Connectivity between SMN 2 and Lateral VN 2 | Mid-life computer use | 0.084 | 0.026 | 0.039 | 0.001 | 0.172 |
| Connectivity between R FPN and BN | Mid-life family and friend visits | -0.056 | 0.017 | -0.038 | 0.001 | 0.172 |
| Connectivity between SMN 3 and Precuneus/PCC DMN | Mid-life participation in sports clubs or gyms | 0.043 | 0.013 | 0.039 | 0.001 | 0.184 |
| Connectivity between Medial VN and Lateral VN 2 | Mid-life attendance to social clubs or pubs | 0.114 | 0.036 | 0.041 | 0.002 | 0.184 |
| Connectivity between SMN 2 and SMN 3 | Mid-life attendance to adult educational classes | 0.110 | 0.036 | 0.037 | 0.002 | 0.197 |
| Connectivity between L FPN and SMN 2 | Mid-life participation in sports clubs or gyms | 0.057 | 0.018 | 0.038 | 0.002 | 0.197 |
| Connectivity between AN and SMN 3 | Mid-life participation in sports clubs or gyms | 0.053 | 0.017 | 0.037 | 0.002 | 0.197 |
| Connectivity between Medial VN and ECN | Mid-life attendance to adult educational classes | 0.071 | 0.024 | 0.036 | 0.003 | 0.217 |
| Connectivity between R FPN and Anterior DMN 2 | Mid-life participation in sports clubs or gyms | -0.054 | 0.018 | -0.037 | 0.003 | 0.218 |
| Connectivity between SMN 2 and SMN 3 | Mid-life participation in sports clubs or gyms | 0.069 | 0.024 | 0.035 | 0.004 | 0.254 |
| Connectivity between ECN and AN 2 | Mid-life participation in sports clubs or gyms | -0.059 | 0.020 | -0.035 | 0.004 | 0.254 |
| Connectivity between Medial VN 2 and SMN 2 | Mid-life participation in sports clubs or gyms | 0.055 | 0.019 | 0.034 | 0.005 | 0.305 |
| Connectivity between ECN and Lateral VN 2 | Mid-life participation in sports clubs or gyms | 0.050 | 0.018 | 0.034 | 0.005 | 0.305 |
| Connectivity between SMN and Medial VN 2 | Mid-life attendance to adult educational classes | -0.085 | 0.030 | -0.034 | 0.005 | 0.305 |
| Connectivity between Anterior DMN and Lateral VN 2 | Mid-life family and friend visits | 0.057 | 0.020 | 0.033 | 0.005 | 0.308 |
| Connectivity between CN and Precuneus/PCC DMN | Mid-life family and friend visits | -0.044 | 0.016 | -0.032 | 0.006 | 0.311 |
| Connectivity between AN and AN 2 | Mid-life computer use | 0.068 | 0.025 | 0.033 | 0.006 | 0.313 |
| Connectivity between SMN 3 and Anterior DMN 2 | Mid-life computer use | -0.050 | 0.018 | -0.033 | 0.006 | 0.313 |
| Connectivity between SMN 2 and CN | Mid-life participation in sports clubs or gyms | 0.048 | 0.018 | 0.033 | 0.007 | 0.313 |
| Connectivity between Anterior DMN 2 and R CON | Mid-life family and friend visits | -0.056 | 0.021 | -0.032 | 0.007 | 0.313 |
| Connectivity between Lateral VN and SMN 2 | Mid-life participation in religious activities | -0.069 | 0.025 | -0.034 | 0.007 | 0.313 |
| Connectivity between SMN 3 and Lateral VN 2 | Mid-life participation in religious activities | 0.044 | 0.016 | 0.034 | 0.007 | 0.313 |
| Connectivity between Anterior DMN and Precuneus/PCC DMN | Mid-life participation in sports clubs or gyms | 0.057 | 0.021 | 0.032 | 0.007 | 0.313 |
| Connectivity between AN and BN | Mid-life family and friend visits | -0.052 | 0.019 | -0.032 | 0.007 | 0.313 |
| Connectivity between CN and AN 2 | Mid-life participation in sports clubs or gyms | 0.040 | 0.015 | 0.032 | 0.007 | 0.313 |

| | | | | | | |
|---|---|---|---|---|---|---|
| Connectivity between R FPN and R CON | Mid-life participation in religious activities | 0.074 | 0.028 | 0.034 | 0.007 | 0.313 |
| Connectivity between SMN 2 and AN | Mid-life participation in religious activities | 0.066 | 0.025 | 0.033 | 0.008 | 0.313 |
| Connectivity between Anterior DMN and R FPN | Mid-life participation in religious activities | 0.074 | 0.028 | 0.033 | 0.009 | 0.343 |
| Connectivity between Anterior DMN and R CON | Mid-life participation in sports clubs or gyms | -0.052 | 0.020 | -0.032 | 0.009 | 0.343 |
| Connectivity between Precuneus/PCC DMN and R CON | Mid-life participation in sports clubs or gyms | -0.048 | 0.019 | -0.032 | 0.010 | 0.343 |
| Connectivity between Posterior DMN and ECN | Mid-life attendance to social clubs or pubs | -0.062 | 0.024 | -0.034 | 0.010 | 0.343 |
| Connectivity between R FPN and SMN 2 | Mid-life attendance to adult educational classes | 0.066 | 0.026 | 0.031 | 0.011 | 0.343 |
| Connectivity between Lateral VN and Posterior DMN | Mid-life attendance to adult educational classes | -0.080 | 0.032 | -0.030 | 0.011 | 0.343 |
| Connectivity between L FPN and ECN | Mid-life family and friend visits | -0.073 | 0.029 | -0.030 | 0.011 | 0.343 |
| Connectivity between Posterior DMN and Lateral VN 2 | Mid-life computer use | 0.063 | 0.025 | 0.030 | 0.012 | 0.343 |
| Connectivity between SMN and SMN 3 | Mid-life participation in sports clubs or gyms | -0.043 | 0.017 | -0.030 | 0.012 | 0.343 |
| Connectivity between Posterior DMN 2 and Lateral VN 2 | Mid-life participation in sports clubs or gyms | -0.044 | 0.017 | -0.031 | 0.012 | 0.345 |
| Connectivity between L CON and Anterior DMN 2 | Mid-life participation in sports clubs or gyms | 0.040 | 0.016 | 0.031 | 0.012 | 0.351 |
| Connectivity between Medial VN 2 and Lateral VN 2 | Mid-life participation in sports clubs or gyms | 0.090 | 0.036 | 0.030 | 0.013 | 0.362 |
| Connectivity between L CON and R CON | Mid-life participation in religious activities | -0.057 | 0.023 | -0.031 | 0.014 | 0.371 |
| Connectivity between AN and BN | Mid-life participation in sports clubs or gyms | -0.039 | 0.016 | -0.030 | 0.014 | 0.372 |
| Connectivity between Medial VN and L CON | Mid-life attendance to adult educational classes | -0.046 | 0.019 | -0.030 | 0.015 | 0.375 |
| Connectivity between Anterior DMN and Lateral VN 2 | Mid-life attendance to adult educational classes | 0.060 | 0.025 | 0.029 | 0.015 | 0.375 |
| Connectivity between Anterior DMN 2 and AN 2 | Mid-life computer use | -0.064 | 0.026 | -0.029 | 0.015 | 0.375 |
| Connectivity between Lateral VN and Anterior DMN 2 | Mid-life participation in sports clubs or gyms | 0.035 | 0.014 | 0.030 | 0.016 | 0.381 |
| Connectivity between AN and Lateral VN 2 | Mid-life participation in sports clubs or gyms | 0.029 | 0.012 | 0.029 | 0.016 | 0.381 |
| Connectivity between SMN 3 and BN | Mid-life family and friend visits | -0.046 | 0.019 | -0.028 | 0.016 | 0.382 |
| Connectivity between Lateral VN and AN 2 | Mid-life participation in sports clubs or gyms | -0.054 | 0.023 | -0.029 | 0.017 | 0.392 |
| Connectivity between Posterior DMN 2 and Precuneus/PCC DMN | Mid-life attendance to social clubs or pubs | -0.062 | 0.026 | -0.031 | 0.017 | 0.392 |
| Connectivity between AN and AN 2 | Mid-life participation in sports clubs or gyms | 0.038 | 0.016 | 0.029 | 0.017 | 0.392 |
| Connectivity between R FPN and AN 2 | Mid-life attendance to social clubs or pubs | 0.057 | 0.024 | 0.031 | 0.018 | 0.395 |
| Connectivity between Lateral VN and Lateral VN 2 | Mid-life attendance to adult educational classes | -0.088 | 0.037 | -0.028 | 0.018 | 0.395 |
| Connectivity between SMN and Posterior DMN | Mid-life computer use | -0.078 | 0.033 | -0.028 | 0.018 | 0.395 |
| Connectivity between SMN 2 and Precuneus/PCC DMN | Mid-life attendance to adult educational classes | 0.059 | 0.025 | 0.028 | 0.019 | 0.399 |
| Connectivity between R FPN and SMN 2 | Mid-life attendance to social clubs or pubs | 0.046 | 0.020 | 0.031 | 0.019 | 0.404 |
| Connectivity between Posterior DMN 2 and Precuneus/PCC DMN | Mid-life participation in religious activities | 0.065 | 0.028 | 0.029 | 0.020 | 0.409 |
| Connectivity between Medial VN and Posterior DMN | Mid-life attendance to adult educational classes | -0.054 | 0.023 | -0.028 | 0.021 | 0.409 |
| Connectivity between SMN and Medial VN 2 | Mid-life attendance to social clubs or pubs | 0.053 | 0.023 | 0.030 | 0.021 | 0.409 |
| Connectivity between Posterior DMN and SMN 2 | Mid-life attendance to adult educational classes | 0.064 | 0.028 | 0.028 | 0.021 | 0.409 |

| | | | | | | |
|---|---|---|---|---|---|---|
| Connectivity between SMN 3 and L CON | Mid-life participation in sports clubs or gyms | -0.028 | 0.012 | -0.028 | 0.021 | 0.409 |
| Connectivity between Medial VN 2 and L CON | Mid-life attendance to social clubs or pubs | -0.041 | 0.018 | -0.030 | 0.022 | 0.416 |
| Connectivity between Anterior DMN 2 and BN | Mid-life family and friend visits | 0.036 | 0.016 | 0.027 | 0.023 | 0.421 |
| Connectivity between SMN 3 and CN | Mid-life attendance to adult educational classes | 0.048 | 0.021 | 0.027 | 0.024 | 0.427 |
| Connectivity between SMN 2 and AN | Mid-life family and friend visits | 0.057 | 0.025 | 0.027 | 0.024 | 0.427 |
| Connectivity between Anterior DMN 2 and ECN | Mid-life participation in sports clubs or gyms | -0.040 | 0.018 | -0.027 | 0.024 | 0.427 |
| Connectivity between CN and BN | Mid-life family and friend visits | 0.032 | 0.014 | 0.026 | 0.024 | 0.427 |
| Connectivity between SMN 2 and Anterior DMN 2 | Mid-life family and friend visits | 0.043 | 0.019 | 0.027 | 0.025 | 0.427 |
| Connectivity between Posterior DMN 2 and AN 2 | Mid-life attendance to social clubs or pubs | 0.051 | 0.023 | 0.029 | 0.025 | 0.427 |
| Connectivity between Lateral VN and AN | Mid-life participation in religious activities | -0.036 | 0.016 | -0.028 | 0.026 | 0.427 |
| Connectivity between Posterior DMN and AN 2 | Mid-life participation in sports clubs or gyms | 0.044 | 0.020 | 0.027 | 0.027 | 0.427 |
| Connectivity between L CON and Precuneus/PCC DMN | Mid-life attendance to adult educational classes | 0.058 | 0.026 | 0.027 | 0.027 | 0.427 |
| Connectivity between Lateral VN and SMN 2 | Mid-life family and friend visits | -0.057 | 0.026 | -0.026 | 0.027 | 0.427 |
| Connectivity between SMN and CN | Mid-life participation in sports clubs or gyms | 0.027 | 0.012 | 0.027 | 0.027 | 0.427 |
| Connectivity between Anterior DMN and BN | Mid-life participation in religious activities | 0.036 | 0.016 | 0.028 | 0.028 | 0.428 |
| Connectivity between Posterior DMN and ECN | Mid-life participation in religious activities | -0.057 | 0.026 | -0.028 | 0.028 | 0.432 |
| Connectivity between Medial VN 2 and SMN 2 | Mid-life participation in religious activities | 0.051 | 0.023 | 0.027 | 0.029 | 0.435 |
| Connectivity between Medial VN 2 and AN 2 | Mid-life participation in sports clubs or gyms | 0.049 | 0.023 | 0.027 | 0.029 | 0.435 |
| Connectivity between R FPN and Posterior DMN | Mid-life participation in sports clubs or gyms | -0.048 | 0.022 | -0.027 | 0.029 | 0.435 |
| Connectivity between L FPN and ECN | Mid-life participation in sports clubs or gyms | -0.051 | 0.023 | -0.026 | 0.030 | 0.442 |
| Connectivity between Lateral VN and AN 2 | Mid-life attendance to social clubs or pubs | 0.055 | 0.025 | 0.028 | 0.030 | 0.446 |
| Connectivity between Medial VN and AN 2 | Mid-life computer use | 0.058 | 0.027 | 0.026 | 0.031 | 0.450 |
| Connectivity between Posterior DMN and SMN 2 | Mid-life participation in sports clubs or gyms | 0.040 | 0.019 | 0.026 | 0.031 | 0.452 |
| Connectivity between Anterior DMN and Precuneus/PCC DMN | Mid-life attendance to social clubs or pubs | 0.051 | 0.024 | 0.027 | 0.032 | 0.458 |
| Connectivity between SMN and ECN | Mid-life attendance to social clubs or pubs | -0.049 | 0.023 | -0.028 | 0.032 | 0.458 |
| Connectivity between CN and R CON | Mid-life participation in religious activities | -0.027 | 0.013 | -0.027 | 0.032 | 0.458 |
| Connectivity between Anterior DMN 2 and ECN | Mid-life computer use | 0.058 | 0.027 | 0.025 | 0.034 | 0.461 |
| Connectivity between AN 2 and Lateral VN 2 | Mid-life family and friend visits | 0.052 | 0.025 | 0.025 | 0.034 | 0.461 |
| Connectivity between L FPN and SMN 3 | Mid-life attendance to social clubs or pubs | 0.034 | 0.016 | 0.028 | 0.034 | 0.461 |
| Connectivity between SMN and SMN 3 | Mid-life computer use | -0.056 | 0.027 | -0.024 | 0.035 | 0.461 |
| Connectivity between L FPN and Posterior DMN | Mid-life computer use | -0.068 | 0.032 | -0.025 | 0.035 | 0.461 |
| Connectivity between Medial VN and Posterior DMN | Mid-life participation in religious activities | -0.040 | 0.019 | -0.026 | 0.035 | 0.461 |
| Connectivity between SMN 3 and R CON | Mid-life attendance to social clubs or pubs | -0.029 | 0.014 | -0.027 | 0.035 | 0.461 |
| Connectivity between Posterior DMN and L CON | Mid-life family and friend visits | -0.050 | 0.024 | -0.025 | 0.035 | 0.461 |

| | | | | | | |
|---|---|---|---|---|---|---|
| Connectivity between Lateral VN and SMN 2 | Mid-life attendance to adult educational classes | -0.066 | 0.031 | -0.025 | 0.035 | 0.461 |
| Connectivity between Medial VN and Medial VN 2 | Mid-life attendance to adult educational classes | -0.083 | 0.039 | -0.025 | 0.036 | 0.461 |
| Connectivity between Posterior DMN 2 and BN | Mid-life attendance to social clubs or pubs | 0.029 | 0.014 | 0.027 | 0.036 | 0.461 |
| Connectivity between Posterior DMN 2 and Medial VN 2 | Mid-life participation in sports clubs or gyms | -0.043 | 0.021 | -0.025 | 0.037 | 0.469 |
| Connectivity between Anterior DMN and BN | Mid-life attendance to adult educational classes | 0.042 | 0.020 | 0.025 | 0.038 | 0.469 |
| Connectivity between Posterior DMN and Precuneus/PCC DMN | Mid-life family and friend visits | -0.055 | 0.026 | -0.024 | 0.039 | 0.469 |
| Connectivity between Posterior DMN 2 and SMN 2 | Mid-life participation in sports clubs or gyms | 0.035 | 0.017 | 0.025 | 0.039 | 0.469 |
| Connectivity between SMN 3 and R CON | Mid-life family and friend visits | 0.031 | 0.015 | 0.024 | 0.039 | 0.469 |
| Connectivity between Medial VN and L FPN | Mid-life attendance to adult educational classes | -0.046 | 0.022 | -0.025 | 0.041 | 0.469 |
| Connectivity between Posterior DMN 2 and Anterior DMN 2 | Mid-life computer use | 0.055 | 0.027 | 0.025 | 0.041 | 0.469 |
| Connectivity between Medial VN 2 and BN | Mid-life family and friend visits | -0.033 | 0.016 | -0.024 | 0.041 | 0.469 |
| Connectivity between Anterior DMN and Lateral VN | Mid-life participation in sports clubs or gyms | -0.036 | 0.017 | -0.025 | 0.041 | 0.469 |
| Connectivity between SMN and ECN | Mid-life participation in religious activities | -0.050 | 0.025 | -0.026 | 0.041 | 0.469 |
| Connectivity between Anterior DMN and Medial VN 2 | Mid-life family and friend visits | 0.045 | 0.022 | 0.024 | 0.041 | 0.469 |
| Connectivity between BN and Precuneus/PCC DMN | Mid-life attendance to adult educational classes | -0.037 | 0.018 | -0.025 | 0.042 | 0.469 |
| Connectivity between R FPN and AN 2 | Mid-life participation in sports clubs or gyms | 0.044 | 0.021 | 0.025 | 0.042 | 0.469 |
| Connectivity between Anterior DMN 2 and BN | Mid-life participation in religious activities | 0.031 | 0.015 | 0.025 | 0.043 | 0.469 |
| Connectivity between AN and R CON | Mid-life attendance to social clubs or pubs | 0.027 | 0.014 | 0.026 | 0.043 | 0.469 |
| Connectivity between SMN 2 and ECN | Mid-life participation in sports clubs or gyms | 0.037 | 0.018 | 0.024 | 0.043 | 0.469 |
| Connectivity between SMN 3 and BN | Mid-life attendance to adult educational classes | -0.047 | 0.023 | -0.024 | 0.044 | 0.469 |
| Connectivity between SMN 2 and ECN | Mid-life participation in religious activities | -0.044 | 0.022 | -0.025 | 0.044 | 0.469 |
| Connectivity between L CON and Anterior DMN 2 | Mid-life attendance to social clubs or pubs | 0.036 | 0.018 | 0.026 | 0.044 | 0.469 |
| Connectivity between Medial VN 2 and SMN 3 | Mid-life participation in sports clubs or gyms | 0.030 | 0.015 | 0.025 | 0.044 | 0.469 |
| Connectivity between SMN 2 and CN | Mid-life attendance to adult educational classes | 0.053 | 0.026 | 0.024 | 0.044 | 0.469 |
| Connectivity between SMN and AN 2 | Mid-life computer use | 0.063 | 0.031 | 0.024 | 0.044 | 0.469 |
| Connectivity between SMN and ECN | Mid-life attendance to adult educational classes | -0.061 | 0.031 | -0.024 | 0.045 | 0.474 |
| Connectivity between SMN and Medial VN | Mid-life participation in religious activities | -0.037 | 0.019 | -0.025 | 0.046 | 0.475 |
| Connectivity between SMN and SMN 3 | Mid-life attendance to adult educational classes | -0.051 | 0.026 | -0.023 | 0.046 | 0.475 |
| Connectivity between R FPN and R CON | Mid-life participation in sports clubs or gyms | -0.046 | 0.023 | -0.024 | 0.046 | 0.478 |
| Connectivity between AN 2 and BN | Mid-life attendance to social clubs or pubs | -0.036 | 0.018 | -0.026 | 0.047 | 0.484 |
| Connectivity between Anterior DMN 2 and CN | Mid-life family and friend visits | -0.026 | 0.013 | -0.023 | 0.048 | 0.484 |
| Connectivity between SMN 2 and ECN | Mid-life family and friend visits | 0.044 | 0.022 | 0.023 | 0.048 | 0.485 |
| Connectivity between Anterior DMN and Posterior DMN | Mid-life participation in religious activities | 0.051 | 0.026 | 0.025 | 0.048 | 0.485 |
| Connectivity between Anterior DMN 2 and CN | Mid-life computer use | -0.033 | 0.017 | -0.023 | 0.049 | 0.485 |

| | | | | | | |
|---|---|---|---|---|---|---|
| Connectivity between SMN 2 and Anterior DMN 2 | Mid-life participation in sports clubs or gyms | 0.030 | 0.015 | 0.024 | 0.050 | 0.492 |

**Abbreviations-** AN = Auditory Network; BGN= Basal ganglia network; CN = Cerebellar Network; CON = Cingulo-opercular network; DMN = Default mode network; ECN = Executive control network; FPN = Frontoparietal Network; SMN = Sensorimotor network; VN = Visual Network.

**Bold = survived FDR corrections.**

**Table S8. Table to show the associations between weekly mid-life participation in activities and resting-state functional connectivity.** Results are adjusted for sociodemographic, imaging, health and lifestyle co-variates.

| Dependent Variable | Predictor | B | SE | β | p-value | FDR q-value |
|---|---|---|---|---|---|---|
| Connectivity between SMN 3 and CN | Late-life participation in sports clubs or gyms | 0.055 | 0.014 | 0.045 | 0.000 | 0.256 |
| Connectivity between SMN 3 and Lateral VN 2 | Late-life participation in sports clubs or gyms | 0.049 | 0.014 | 0.043 | 0.000 | 0.256 |
| Connectivity between R FPN and SMN 3 | Late-life computer use | -0.110 | 0.031 | -0.042 | 0.000 | 0.256 |
| Connectivity between AN and AN 2 | Late-life participation in sports clubs or gyms | 0.055 | 0.016 | 0.042 | 0.001 | 0.256 |
| Connectivity between AN and CN | Late-life participation in sports clubs or gyms | 0.044 | 0.013 | 0.041 | 0.001 | 0.256 |
| Connectivity between L FPN and ECN | Late-life participation in sports clubs or gyms | -0.078 | 0.023 | -0.041 | 0.001 | 0.256 |
| Connectivity between Anterior DMN and BN | Late-life participation in religious activities | 0.052 | 0.016 | 0.040 | 0.001 | 0.337 |
| Connectivity between Anterior DMN and Precuneus/PCC DMN | Late-life participation in sports clubs or gyms | 0.068 | 0.021 | 0.038 | 0.001 | 0.345 |
| Connectivity between SMN 2 and SMN 3 | Late-life participation in sports clubs or gyms | 0.074 | 0.024 | 0.037 | 0.002 | 0.345 |
| Connectivity between Lateral VN and SMN 2 | Late-life participation in religious activities | -0.076 | 0.025 | -0.038 | 0.003 | 0.345 |
| Connectivity between AN and SMN 3 | Late-life participation in sports clubs or gyms | 0.052 | 0.017 | 0.036 | 0.003 | 0.345 |
| Connectivity between SMN 2 and Anterior DMN 2 | Late-life participation in sports clubs or gyms | 0.046 | 0.015 | 0.036 | 0.003 | 0.345 |
| Connectivity between SMN 2 and ECN | Late-life attendance to adult educational classes | 0.085 | 0.029 | 0.035 | 0.003 | 0.345 |
| Connectivity between SMN and SMN 3 | Late-life participation in sports clubs or gyms | -0.050 | 0.017 | -0.035 | 0.003 | 0.345 |
| Connectivity between Anterior DMN 2 and BN | Late-life participation in religious activities | 0.044 | 0.015 | 0.036 | 0.004 | 0.345 |
| Connectivity between Medial VN and R FPN | Late-life family and friend visits | -0.059 | 0.020 | -0.035 | 0.004 | 0.345 |
| Connectivity between Lateral VN and AN | Late-life participation in religious activities | -0.046 | 0.016 | -0.036 | 0.004 | 0.345 |
| Connectivity between Medial VN 2 and Posterior DMN | Late-life participation in sports clubs or gyms | 0.053 | 0.019 | 0.035 | 0.004 | 0.345 |
| Connectivity between L CON and R CON | Late-life attendance to adult educational classes | -0.085 | 0.030 | -0.034 | 0.005 | 0.345 |
| Connectivity between SMN 2 and R CON | Late-life participation in sports clubs or gyms | 0.047 | 0.017 | 0.034 | 0.005 | 0.345 |
| Connectivity between SMN and Posterior DMN | Late-life attendance to social clubs or pubs | 0.067 | 0.024 | 0.036 | 0.005 | 0.345 |
| Connectivity between Posterior DMN 2 and Medial VN 2 | Late-life computer use | 0.125 | 0.045 | 0.033 | 0.006 | 0.345 |
| Connectivity between SMN 3 and Lateral VN 2 | Late-life participation in religious activities | 0.045 | 0.016 | 0.035 | 0.006 | 0.345 |
| Connectivity between Medial VN and R CON | Late-life attendance to social clubs or pubs | 0.039 | 0.014 | 0.036 | 0.006 | 0.345 |
| Connectivity between SMN 3 and BN | Late-life participation in sports clubs or gyms | -0.043 | 0.015 | -0.033 | 0.006 | 0.345 |
| Connectivity between CN and AN 2 | Late-life attendance to adult educational classes | 0.065 | 0.024 | 0.033 | 0.006 | 0.345 |
| Connectivity between AN and Lateral VN 2 | Late-life attendance to social clubs or pubs | -0.037 | 0.013 | -0.035 | 0.006 | 0.345 |
| Connectivity between CN and ECN | Late-life family and friend visits | 0.046 | 0.017 | 0.032 | 0.006 | 0.345 |
| Connectivity between CN and Precuneus/PCC DMN | Late-life attendance to social clubs or pubs | 0.039 | 0.014 | 0.034 | 0.006 | 0.352 |
| Connectivity between R FPN and AN 2 | Late-life attendance to social clubs or pubs | 0.064 | 0.024 | 0.035 | 0.007 | 0.357 |
| Connectivity between Anterior DMN and Lateral VN | Late-life participation in sports clubs or gyms | -0.047 | 0.017 | -0.033 | 0.008 | 0.357 |
| Connectivity between Anterior DMN and AN | Late-life attendance to adult educational classes | 0.055 | 0.021 | 0.032 | 0.008 | 0.357 |
| Connectivity between L CON and AN 2 | Late-life participation in religious activities | -0.062 | 0.023 | -0.033 | 0.008 | 0.357 |

| | | | | | | |
|---|---|---|---|---|---|---|
| Connectivity between SMN 2 and AN | Late-life participation in religious activities | 0.065 | 0.024 | 0.033 | 0.008 | 0.357 |
| Connectivity between L FPN andPrecuneus/PCC DMN | Late-life attendance to adult educational classes | 0.090 | 0.034 | 0.032 | 0.008 | 0.357 |
| Connectivity between SMN and R CON | Late-life participation in sports clubs or gyms | -0.051 | 0.019 | -0.032 | 0.008 | 0.357 |
| Connectivity between Medial VN and Posterior DMN | Late-life participation in religious activities | -0.050 | 0.019 | -0.033 | 0.008 | 0.357 |
| Connectivity between L CON and Anterior DMN 2 | Late-life participation in sports clubs or gyms | 0.042 | 0.016 | 0.032 | 0.008 | 0.357 |
| Connectivity between Anterior DMN 2 and AN 2 | Late-life participation in sports clubs or gyms | 0.044 | 0.017 | 0.032 | 0.008 | 0.357 |
| Connectivity between AN and AN 2 | Late-life computer use | 0.093 | 0.035 | 0.031 | 0.009 | 0.357 |
| Connectivity between Medial VN and Posterior DMN | Late-life attendance to adult educational classes | -0.064 | 0.025 | -0.031 | 0.009 | 0.357 |
| Connectivity between Medial VN 2 and Posterior DMN | Late-life computer use | 0.106 | 0.041 | 0.031 | 0.009 | 0.357 |
| Connectivity between Posterior DMN and R CON | Late-life family and friend visits | 0.064 | 0.025 | 0.031 | 0.010 | 0.363 |
| Connectivity between Medial VN and L FPN | Late-life attendance to adult educational classes | -0.061 | 0.024 | -0.031 | 0.011 | 0.363 |
| Connectivity between Anterior DMN 2 and CN | Late-life family and friend visits | -0.035 | 0.014 | -0.030 | 0.011 | 0.363 |
| Connectivity between R FPN and SMN 2 | Late-life computer use | 0.097 | 0.038 | 0.030 | 0.011 | 0.365 |
| Connectivity between AN 2 and Lateral VN 2 | Late-life participation in sports clubs or gyms | 0.051 | 0.020 | 0.031 | 0.012 | 0.369 |
| Connectivity between L CON and Anterior DMN 2 | Late-life computer use | -0.089 | 0.035 | -0.030 | 0.012 | 0.369 |
| Connectivity between Lateral VN and CN | Late-life family and friend visits | 0.041 | 0.016 | 0.030 | 0.012 | 0.369 |
| Connectivity between Lateral VN and SMN 2 | Late-life attendance to adult educational classes | -0.084 | 0.033 | -0.030 | 0.012 | 0.369 |
| Connectivity between AN 2 andPrecuneus/PCC DMN | Late-life computer use | -0.100 | 0.040 | -0.030 | 0.012 | 0.369 |
| Connectivity between CN and AN 2 | Late-life participation in sports clubs or gyms | 0.037 | 0.015 | 0.030 | 0.012 | 0.369 |
| Connectivity between Lateral VN and AN 2 | Late-life participation in sports clubs or gyms | -0.056 | 0.023 | -0.029 | 0.014 | 0.410 |
| Connectivity between SMN and CN | Late-life participation in sports clubs or gyms | 0.030 | 0.012 | 0.029 | 0.017 | 0.455 |
| Connectivity between Medial VN 2 and Posterior DMN | Late-life family and friend visits | 0.056 | 0.024 | 0.028 | 0.017 | 0.463 |
| Connectivity between SMN and Anterior DMN 2 | Late-life attendance to social clubs or pubs | 0.042 | 0.018 | 0.030 | 0.018 | 0.483 |
| Connectivity between ECN andPrecuneus/PCC DMN | Late-life family and friend visits | 0.059 | 0.025 | 0.027 | 0.020 | 0.510 |
| Connectivity between Posterior DMN 2 and Medial VN 2 | Late-life participation in sports clubs or gyms | -0.047 | 0.021 | -0.028 | 0.021 | 0.517 |
| Connectivity between Medial VN and SMN 2 | Late-life attendance to adult educational classes | -0.058 | 0.026 | -0.028 | 0.022 | 0.538 |
| Connectivity between SMN andPrecuneus/PCC DMN | Late-life attendance to social clubs or pubs | 0.050 | 0.022 | 0.029 | 0.023 | 0.542 |
| Connectivity between SMN 3 and Anterior DMN 2 | Late-life computer use | -0.059 | 0.026 | -0.027 | 0.023 | 0.542 |
| Connectivity between Posterior DMN 2 and Lateral VN 2 | Late-life participation in sports clubs or gyms | -0.039 | 0.017 | -0.028 | 0.024 | 0.545 |
| Connectivity between R FPN and R CON | Late-life attendance to social clubs or pubs | -0.057 | 0.025 | -0.029 | 0.024 | 0.545 |
| Connectivity between Posterior DMN and SMN 3 | Late-life attendance to social clubs or pubs | -0.032 | 0.014 | -0.029 | 0.026 | 0.562 |
| Connectivity between Lateral VN and R FPN | Late-life attendance to adult educational classes | 0.066 | 0.030 | 0.027 | 0.026 | 0.562 |
| Connectivity between Posterior DMN 2 and ECN | Late-life participation in sports clubs or gyms | -0.050 | 0.023 | -0.027 | 0.027 | 0.564 |
| Connectivity between SMN and Posterior DMN | Late-life computer use | -0.111 | 0.050 | -0.026 | 0.027 | 0.564 |

| | | | | | | |
|---|---|---|---|---|---|---|
| Connectivity between Anterior DMN and Lateral VN | Late-life attendance to social clubs or pubs | 0.043 | 0.019 | 0.028 | 0.027 | 0.564 |
| Connectivity between L CON and BN | Late-life participation in religious activities | -0.031 | 0.014 | -0.028 | 0.028 | 0.564 |
| Connectivity between SMN 3 and AN 2 | Late-life participation in sports clubs or gyms | 0.038 | 0.017 | 0.027 | 0.028 | 0.566 |
| Connectivity between Medial VN 2 and AN 2 | Late-life computer use | 0.108 | 0.050 | 0.026 | 0.030 | 0.601 |
| Connectivity between SMN 3 and L CON | Late-life participation in sports clubs or gyms | -0.026 | 0.012 | -0.026 | 0.031 | 0.602 |
| Connectivity between Posterior DMN and AN 2 | Late-life participation in sports clubs or gyms | 0.043 | 0.020 | 0.026 | 0.031 | 0.602 |
| Connectivity between Anterior DMN and CN | Late-life attendance to adult educational classes | -0.044 | 0.021 | -0.026 | 0.032 | 0.602 |
| Connectivity between SMN 3 and Precuneus/PCC DMN | Late-life attendance to adult educational classes | 0.046 | 0.021 | 0.026 | 0.034 | 0.623 |
| Connectivity between AN and AN 2 | Late-life participation in religious activities | 0.041 | 0.019 | 0.026 | 0.034 | 0.624 |
| Connectivity between Medial VN 2 and SMN 3 | Late-life participation in sports clubs or gyms | 0.032 | 0.015 | 0.026 | 0.035 | 0.624 |
| Connectivity between Medial VN 2 and L CON | Late-life attendance to social clubs or pubs | -0.037 | 0.018 | -0.027 | 0.035 | 0.624 |
| Connectivity between Anterior DMN 2 and Precuneus/PCC DMN | Late-life computer use | 0.083 | 0.040 | 0.025 | 0.036 | 0.624 |
| Connectivity between L FPN and R CON | Late-life attendance to adult educational classes | 0.091 | 0.044 | 0.025 | 0.036 | 0.624 |
| Connectivity between AN and Lateral VN 2 | Late-life computer use | -0.055 | 0.027 | -0.025 | 0.038 | 0.646 |
| Connectivity between R FPN and Medial VN 2 | Late-life attendance to social clubs or pubs | 0.046 | 0.022 | 0.027 | 0.038 | 0.651 |
| Connectivity between R FPN and ECN | Late-life participation in sports clubs or gyms | 0.047 | 0.023 | 0.025 | 0.040 | 0.652 |
| Connectivity between R FPN and CN | Late-life attendance to adult educational classes | -0.041 | 0.020 | -0.025 | 0.042 | 0.666 |
| Connectivity between L CON and Precuneus/PCC DMN | Late-life computer use | -0.079 | 0.039 | -0.024 | 0.042 | 0.666 |
| Connectivity between R FPN and L CON | Late-life attendance to social clubs or pubs | 0.052 | 0.025 | 0.026 | 0.042 | 0.666 |
| Connectivity between Anterior DMN and CN | Late-life computer use | -0.058 | 0.028 | -0.024 | 0.043 | 0.674 |
| Connectivity between SMN 2 and CN | Late-life attendance to adult educational classes | 0.057 | 0.028 | 0.024 | 0.044 | 0.674 |
| Connectivity between AN and Precuneus/PCC DMN | Late-life computer use | 0.055 | 0.027 | 0.024 | 0.044 | 0.674 |
| Connectivity between Medial VN 2 and BN | Late-life participation in religious activities | -0.031 | 0.016 | -0.025 | 0.046 | 0.698 |
| Connectivity between Posterior DMN and SMN 2 | Late-life computer use | -0.081 | 0.041 | -0.024 | 0.047 | 0.701 |
| Connectivity between Posterior DMN 2 and Medial VN 2 | Late-life participation in religious activities | -0.049 | 0.025 | -0.025 | 0.048 | 0.701 |
| Connectivity between AN and Lateral VN 2 | Late-life participation in sports clubs or gyms | 0.024 | 0.012 | 0.024 | 0.048 | 0.701 |
| Connectivity between SMN and ECN | Late-life attendance to social clubs or pubs | -0.045 | 0.023 | -0.025 | 0.048 | 0.701 |
| Connectivity between AN and CN | Late-life participation in religious activities | 0.031 | 0.016 | 0.025 | 0.048 | 0.701 |
| Connectivity between Posterior DMN 2 and AN 2 | Late-life attendance to adult educational classes | -0.064 | 0.032 | -0.024 | 0.049 | 0.701 |
| Connectivity between R FPN and Anterior DMN 2 | Late-life participation in sports clubs or gyms | -0.035 | 0.018 | -0.024 | 0.049 | 0.701 |
| Connectivity between R FPN and R CON | Late-life participation in religious activities | 0.054 | 0.027 | 0.025 | 0.049 | 0.701 |

**Abbreviations-** AN = Auditory Network; BGN= Basal ganglia network; CN = Cerebellar Network; CON = Cingulo-opercular network; DMN = Default mode network; ECN = Executive control network; FPN = Frontoparietal Network; SMN = Sensorimotor network; VN = Visual Network.

**Bold = survived FDR corrections.**

**Supplementary References**


[1] Fawns-Ritchie C, Deary I. Reliability and validity of the UK Biobank cognitive tests 2019. https://doi.org/10.1101/19002204.

[2] Chan D, Shafto M, Kievit R, Matthews F, Spink M, Valenzuela M, et al. Lifestyle activities in mid-life contribute to cognitive reserve in late-life, independent of education, occupation, and late-life activities. Neurobiol Aging 2018;70:180–3. https://doi.org/10.1016/J.NEUROBIOLAGING.2018.06.012.

[3] Cornelis MC, Wang Y, Holland T, Agarwal P, Weintraub S, Morris MC. Age and cognitive decline in the UK Biobank. PLoS One 2019;14:e0213948. https://doi.org/10.1371/journal.pone.0213948.

[4] Hagenaars SP, Cox SR, Hill WD, Davies G, Liewald DCM, Harris SE, et al. Genetic contributions to Trail Making Test performance in UK Biobank. Mol Psychiatry 2018;23:1575–83. https://doi.org/10.1038/mp.2017.189.

[5] Wu P, Gifford A, Meng X, Li X, Campbell H, Varley T, et al. Developing and Evaluating Mappings of ICD-10 and ICD-10-CM Codes to PheCodes. BioRxiv 2019:462077. https://doi.org/10.1101/462077.

[6] Shankar A, Hamer M, McMunn A, Steptoe A. Social Isolation and Loneliness. Psychosom Med 2013;75:161–70. https://doi.org/10.1097/PSY.0b013e31827f09cd.

[7] Miller KL, Alfaro-Almagro F, Bangerter NK, Thomas DL, Yacoub E, Xu J, et al. Multimodal population brain imaging in the UK Biobank prospective epidemiological study. Nat Neurosci 2016;19:1523–36. https://doi.org/10.1038/nn.4393.

[8] Alfaro-Almagro F, Jenkinson M, Bangerter NK, Andersson JLR, Griffanti L, Douaud G, et al. Image processing and Quality Control for the first 10,000 brain imaging datasets from UK Biobank. Neuroimage 2018;166:400–24. https://doi.org/10.1016/J.NEUROIMAGE.2017.10.034.

[9] Smith SM, Alfaro-Almagro F, Miller KL. UK Biobank Brain Imaging Documentation. 2019.

[10] Smith SM. Fast robust automated brain extraction. Hum Brain Mapp

2002;17:143–55. https://doi.org/10.1002/hbm.10062.
[11] Jenkinson M, Smith S. A global optimisation method for robust affine registration of brain images. Med Image Anal 2001;5:143–56.
[12] Jenkinson M, Bannister P, Brady M, Smith S. Improved optimization for the robust and accurate linear registration and motion correction of brain images. Neuroimage 2002;17:825–41.
[13] Andersson JLR, Jenkinson M, Smith S, Jenkinson M, SMITH S, Andersson JLR, et al. Non-Linear Registration aka Spatial Normalisation FMRIB Technial Report TR07JA2. 2007.
[14] Zhang Y, Brady M, Smith S. Segmentation of brain MR images through a hidden Markov random field model and the expectation-maximization algorithm. IEEE Trans Med Imaging 2001;20:45–57. https://doi.org/10.1109/42.906424.
[15] Patenaude B, Smith SM, Kennedy DN, Jenkinson M. A Bayesian model of shape and appearance for subcortical brain segmentation. Neuroimage 2011;56:907–22. https://doi.org/10.1016/j.neuroimage.2011.02.046.
[16] Smith SM, Zhang Y, Jenkinson M, Chen J, Matthews PM, Federico A, et al. Accurate, robust, and automated longitudinal and cross-sectional brain change analysis. Neuroimage 2002;17:479–89.
[17] Andersson JLR, Skare S, Ashburner J. How to correct susceptibility distortions in spin-echo echo-planar images: application to diffusion tensor imaging. Neuroimage 2003;20:870–88. https://doi.org/10.1016/S1053-8119(03)00336-7.
[18] Andersson JLR, Sotiropoulos SN. Non-parametric representation and prediction of single- and multi-shell diffusion-weighted MRI data using Gaussian processes. Neuroimage 2015;122:166–76. https://doi.org/10.1016/j.neuroimage.2015.07.067.
[19] Andersson JLR, Sotiropoulos SN. An integrated approach to correction for off-resonance effects and subject movement in diffusion MR imaging. Neuroimage 2016;125:1063–78. https://doi.org/10.1016/j.neuroimage.2015.10.019.
[20] Basser PJ, Mattiello J, LeBihan D. MR diffusion tensor spectroscopy and imaging. Biophys J 1994;66:259–67. https://doi.org/10.1016/S0006-

3495(94)80775-1.
[21] Behrens TEJ, Woolrich MW, Jenkinson M, Johansen-Berg H, Nunes RG, Clare S, et al. Characterization and propagation of uncertainty in diffusion-weighted MR imaging. Magn Reson Med 2003;50:1077–88. https://doi.org/10.1002/mrm.10609.
[22] Behrens TEJ, Berg HJ, Jbabdi S, Rushworth MFS, Woolrich MW. Probabilistic diffusion tractography with multiple fibre orientations: What can we gain? Neuroimage 2007;34:144–55. https://doi.org/10.1016/J.NEUROIMAGE.2006.09.018.
[23] Jbabdi S, Sotiropoulos SN, Savio AM, Graña M, Behrens TEJ. Model-based analysis of multishell diffusion MR data for tractography: How to get over fitting problems. Magn Reson Med 2012;68:1846–55. https://doi.org/10.1002/mrm.24204.
[24] Hernández M, Guerrero GD, Cecilia JM, García JM, Inuggi A, Jbabdi S, et al. Accelerating Fibre Orientation Estimation from Diffusion Weighted Magnetic Resonance Imaging Using GPUs. PLoS One 2013;8:e61892. https://doi.org/10.1371/journal.pone.0061892.
[25] de Groot M, Vernooij MW, Klein S, Ikram MA, Vos FM, Smith SM, et al. Improving alignment in Tract-based spatial statistics: Evaluation and optimization of image registration. Neuroimage 2013;76:400–11. https://doi.org/10.1016/j.neuroimage.2013.03.015.
[26] Greve DN, Fischl B. Accurate and robust brain image alignment using boundary-based registration. Neuroimage 2009;48:63–72. https://doi.org/10.1016/j.neuroimage.2009.06.060.
[27] Griffanti L, Zamboni G, Khan A, Li L, Bonifacio G, Sundaresan V, et al. BIANCA (Brain Intensity AbNormality Classification Algorithm): A new tool for automated segmentation of white matter hyperintensities. Neuroimage 2016;141:191–205. https://doi.org/10.1016/j.neuroimage.2016.07.018.
[28] Beckmann CF, Smith SM. Probabilistic Independent Component Analysis for Functional Magnetic Resonance Imaging. IEEE Trans Med Imaging 2004;23:137–52. https://doi.org/10.1109/TMI.2003.822821.
[29] Salimi-Khorshidi G, Douaud G, Beckmann CF, Glasser MF, Griffanti L, Smith SM. Automatic denoising of functional MRI data: Combining

independent component analysis and hierarchical fusion of classifiers. Neuroimage 2014;90:449–68. https://doi.org/10.1016/j.neuroimage.2013.11.046.

[30] Griffanti L, Salimi-Khorshidi G, Beckmann CF, Auerbach EJ, Douaud G, Sexton CE, et al. ICA-based artefact removal and accelerated fMRI acquisition for improved resting state network imaging. Neuroimage 2014;95:232–47.

[31] Shen X, Cox SR, Adams MJ, Howard DM, Lawrie SM, Ritchie SJ, et al. Resting-State Connectivity and Its Association With Cognitive Performance, Educational Attainment, and Household Income in the UK Biobank. Biol Psychiatry Cogn Neurosci Neuroimaging 2018;3:878–86. https://doi.org/10.1016/J.BPSC.2018.06.007.

[32] Smith RX, Jann K, Ances B, Wang DJJ. Wavelet-based regularity analysis reveals recurrent spatiotemporal behavior in resting-state fMRI. Hum Brain Mapp 2015;36:3603–20.
